# Distinct Activation Mechanisms Regulate Subtype Selectivity of Cannabinoid Receptors

**DOI:** 10.1101/2022.09.27.509760

**Authors:** Soumajit Dutta, Diwakar Shukla

## Abstract

Cannabinoid receptors (CB_1_ and CB_2_) are important drug targets for inflammation, obesity, and other central nervous system disorders. However, due to sequence and structural similarities of the ligand binding pockets of these receptors, most of the ligands lack subtype selectivity and cause off-target side effects. CB_2_ selective agonists can potentially treat pain and inflammation without the psychoactive effects of CB_1_ agonism. We hypothesize that the subtype selectivity of designed selective ligands can be explained by ligand binding to the conformationally distinct states between CB_1_ and CB_2_. To find these conformationally distinct states, we perform ∼ 700*μ*s of unbiased simulations to study the activation mechanism of both the receptors in absence of ligands. The simulation datasets of two receptors were analyzed using Markov state models to identify similarities and distinctions of the major conformational changes associated with activation and allosteric communication between them. Specifically, toggle switch residue movement and its effect on receptor activation differ greatly between CB_1_ and CB_2_. Upon further analysis, we discretize the conformational ensembles of both receptors into metastable states using the neural network-based VAMPnets. Structural and dynamic comparisons of these metastable states allow us to decipher a coarse-grained view of protein activation by revealing sequential conversion between these states. Specifically, we observe the difference in the binding pocket volume of different metastable states of CB_1_, whereas there are minimal changes observed in the CB_2_. Docking analysis reveals that differential binding pocket volume leads to distinct binding poses and docking affinities of CB_2_ selective agonists in CB_1_. Only a few of the intermediate metastable states of CB_1_ shows high affinity towards CB_2_ selective agonists. On the other hand, all the CB_2_ metastable states show a similar affinity for CB_2_ selective agonists, explaining these ligands’ overall higher affinity towards CB_2_. Overall, this computational study mechanistically explains the subtype selectivity of CB_2_ selective ligands by deciphering the activation mechanism of cannabinoid receptors.

## Introduction

Cannabinoid receptors (CB_1_ and CB_2_) were first discovered as a target of phytocannabinoids in the last decades of the twentieth-century.^1,2^ Important physiological and psychological roles of these receptors were soon perceived by the elucidation of the endocannabinoid signaling system.^3–5^ CB_1_ is majorly expressed in the central nervous system and has a role in appetite control, neuroprotection, and neurotransmission.^6,7^ CB_2_ is majorly expressed in the immune system; targeted for immunomodulation and inflammation.^6,8^ These receptors belong to the lipid subfamily of class A GPCRs.^9–17^ Class A GPCRs are recognized by conserved motifs which undergo structural changes during activation of the protein and help to transduce intracellular signaling by *β*-arrestins, and G-proteins.^18–20^ GPCR are targets of more than 30% of FDA-approved drugs.^21–23^ Similarly, academic labs and pharmaceutical companies have led drug discovery efforts to develop molecules targeting cannabinoid receptors.^24–27^ Earlier discovery campaigns resulted in molecules structurally similar to the known phytocannabinoids and endocannabinoids, which was followed by generation of chemically diverse molecules.^28^ Most of these ligands bind to both the receptors and lead to nonspecific side effects;^29–31^ only a few designed ligands are selective to CB_1_ or CB_2_.^32^ However, an explanation of mechanistic underpinning for these ligands’ selectivity is still missing. Understanding subtype selectivity between CB_1_ and CB_2_ would be important for selective drug design. Specifically, selective CB_2_ selective agonists could be an important drug target for pain and inflammation without the psychoactive side effects caused by the binding to CB_1_.^33–35^

Lack of subtype selectivity is also a major issue in drug design for other subfamilies of class A GPCRs. For example, the main targets of ergotamine are 5-HT_1*B*_ and 5-HT_1*D*_ receptors; however, its usage is very limited due to off-target actions on the other 5-HT subtype receptors, particularly on the 5-HT_2*B*_ receptor that causes life-threatening cardiovascular disorders.^36^ Structure-based approaches like docking studies or data-driven methods like machine learning have been employed to tackle the subtype selective drug design for class A GPCRs.^37–41^ Large-scale docking studies have been performed where ligands were docked into the static crystal structure or homology models to explain specific interactions crucial for subtype selectivity. For instance, important protein residue difference in TM5 of 5-HT_1*B*_ and 5-HT_2*B*_ was found to be an essential factor in drug design.^42^ However, docking does not take into account the large structural changes in the receptor binding pocket. Therefore, docking might not reveal the correct binding poses for ligands, especially when the binding pocket undergoes a significant volume change. Previous crystal structure studies show that selectivity between A_1_-AR and A_2*A*_-AR appears due to structural rather than sequence changes.^43^ On the contrary, machine learning models (e.g., LiCABEDS) have provided interpretability about the ligand scaffold that might lead to subtype selectivity. However, these ML models fail to consider receptor-ligand interactions, which leads to high false positive rate in their predictions.^44,45^

In the last few years, crystal structures of inactive and active states of CB_1_ and CB_2_ were determined, which revealed ligand binding poses, and important structural changes during the activation of CB_1_ and CB_2_ (Table S1). Comparing the antagonist-bound pose of inactive CB_1_ and CB_2_ shows the distinction between binding poses of antagonist of CB_1_ and CB_2_ (Figure S1A). However, structural comparison of agonist bound pose of unselective ligands of CB_1_ and CB_2_ shows that binding pocket residues have high structural and sequence similarity (Figure S1B and S1E). Docking performed using the experimentally determined structures did not reveal differences in the binding poses nor could determine subtype selectivity for CB_1_ and CB_2_.^9,11^ Furthermore, during the activation process, the CB_1_ binding pocket undergoes large conformational changes due to the upward movement of the N-terminus, whereas CB_2_ pocket retains its overall shape (Figure S1C and S1D). Considering these two factors, we hypothesize that CB_2_ selective agonists conformationally select for the CB_2_ pocket shape. As the pocket shape remains similar through out the activation process, these ligands have the higher possibility of binding to the CB_2_ rather than CB_1_, where the pocket shape and volume changes significantly during the activation thereby decreasing the overall binding affinity (Figure 1).

**Figure 1:**
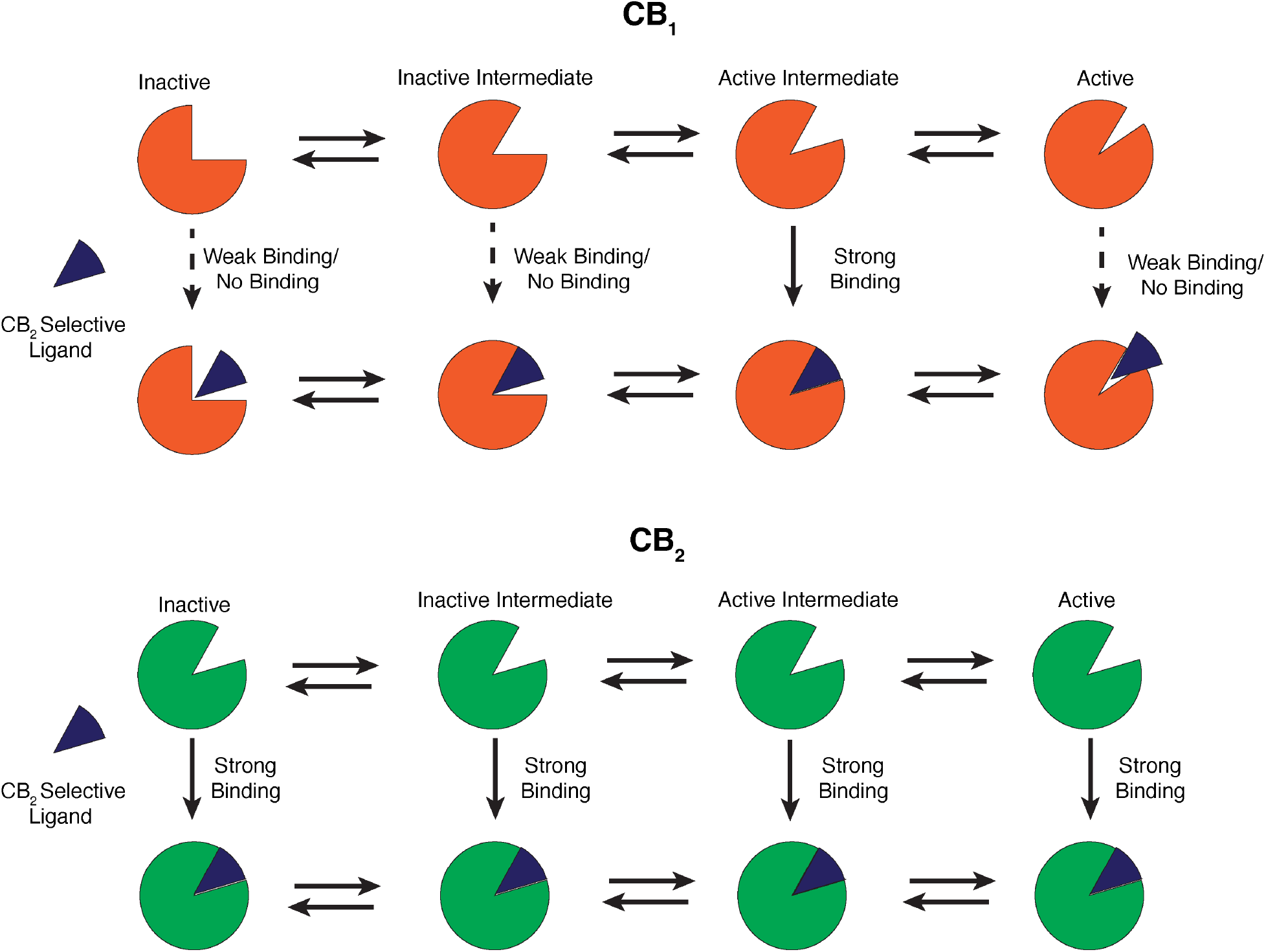
Cartoon representation of the hypothesis describing the subtype selectivity of the CB_2_ selective agonists (Color : Blue). According to the hypothesis, binding pocket volume of CB_1_ (Color: Orange) changes during the activation process. CB_2_ selective agonists may bind into specific intermediate states of CB_1_ decreasing overall binding affnity. Contrastingly, these ligands prefer binding pocket shape of CB_2_ (Color: Green), which retains this generalized shape during activation process.

Here, we compared the activation mechanism for CB_1_ and CB_2_ using molecular dynamics simulations to obtain their entire conformational ensemble. Using 700 *μ*s of aggregate unbiased simulation data, we deciphered the similarity and distinction of activation mechanism between CB_1_ and CB_2_. Analyzing the data using markov state model, we found important allosteric communications between different structural motifs, which facilitates the receptor activation. By implementing deep learning based VAMPnets architecture, we discretize the conformational space into different metastable states.^46^ Docking of the CB_2_ selective ligands shows the docking affinities of these ligands are similar for all the metastable states in CB_2_ whereas these ligands bind to only specific metastable states in CB_1_. Binding of CB_2_ selective ligands favorably to only specific metastable states might explain the subtype selectivity of the ligands. This mechanistic understanding of ligand selectivity between CB_1_ and CB_2_ will aid in the design of new subtype-selective drugs for cannabinoid receptors.

## Results and Discussion

### Structural comparison of CB_1_ and CB_2_ crystal structures

In the last few years, a plethora of CB_1_ and CB_2_ structures are reported in different conditions as shown in Table S1. A comparison of these crystal structures shows major structural changes in the receptors’ extracellular, transmembrane, and intracellular regions during the activation. As both these receptors belong to the lipid subfamily of class A GPCRs, some of these changes are conserved between CB_1_, and CB_2_.^9,11,12,16^ For example, both the receptors undergo large conformational changes in intracellular TM6 towards the outward direction, as shown in Figure 2. Projection of intracellular TM6 movement (R^3.50^-K^6.35^ distance) feature on one dimensional space shows that this feature can distinguish active and inactive structures for both CB_1_ and CB_2_ (Figure 3 and S2). Similarly, the sidechain of conserved Y^7.53^ in the NPxxY region of intracellular TM7 moves towards TM5 during the full activation of the receptors (Figure 2). Projection of intracellular TM7 distance (I^5.54^-Y^7.53^ distance) also shows that this feature changes during activation of CB_1_ and CB_2_ (Figure 3). Therefore, these two conserved intracellular features are used as metrices to judge whether a structure is active or inactive. However, other conformational changes are not conserved between the CB_1_ and CB_2_. Structural overlap of a representative active and inactive CB_1_ and CB_2_ structures show the following non-conserved changes in CB_1_ and CB_2_.

**Figure 2:**
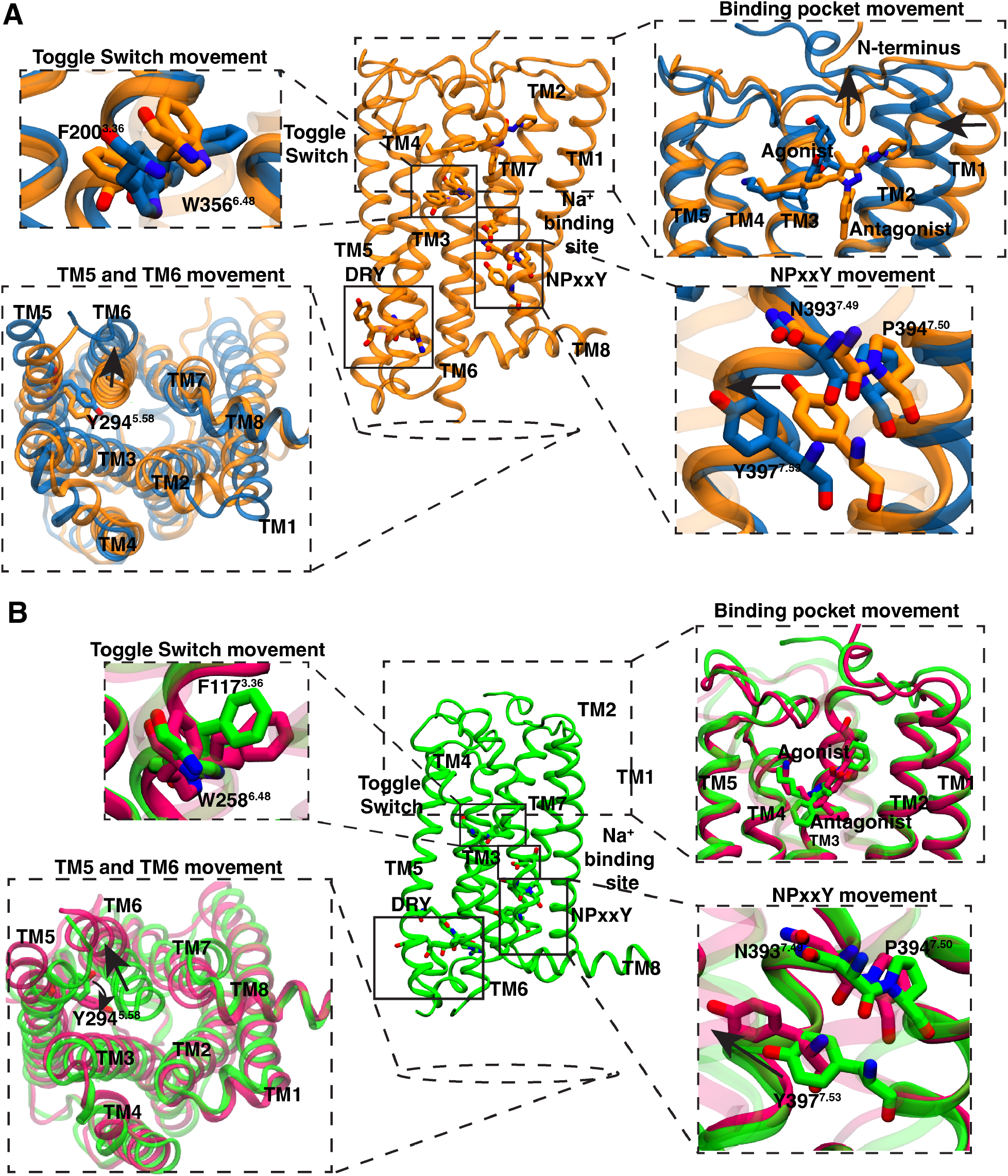
Inactive crystal structure of CB_1_ (A) (PDB ID: 5TGZ,_9_ color: Orange) and CB_2_ (B) (PDB ID: 5TZY_12_, color: Green) are shown as cartoon representations in the central panel. Protein residues in conserved motifs are shown as sticks and highlighted inside block boxes. Major conformational changes during the activation are shown as separated box, where the active (CB_1_ PDB ID: 5XRA,_11_ color: Blue; CB_2_ PDB ID: 6KPF,_16_ color: Pink) and inactive conformations are superposed. Directions of the conformational changes during the activation are shown as black arrow.

**Figure 3:**
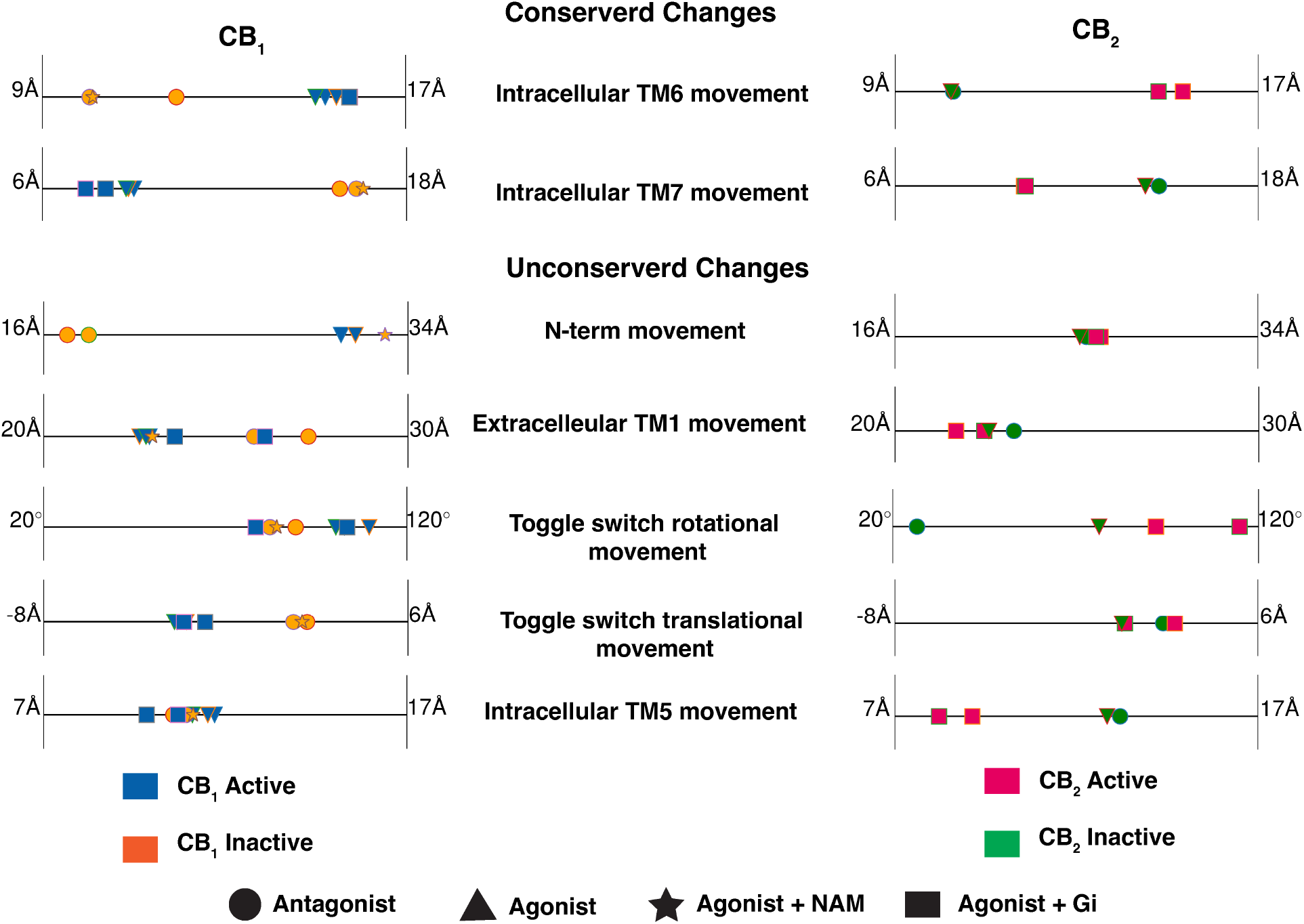
Distance and angle feature values that undergo conformational changes during the activation for CB_1_ (left panel) and CB_2_ (right panel) are shown as scatter points on 1-D line. Markers are colored based on the activation state of the protein. The shapes of the markers are based on the type of the ligand and downstream protein. PDB structures of a receptor with same binding partner and activation state are distinguished with different outline color of the marker.

1. For CB_1_, in the extracellular region, N-terminus shows large conformational differences in agonist bound structure as compared to the antagonist bound structures (Figure 2). Agonist and antagonist bound poses are also different for CB_1_.^11,13^ Agonist binding lifts the N-terminus above the orthosteric pocket. The N-terminus distance (M^N−term^-D^2.50^ distance) from the ligand binding pocket clearly shows the difference in the agonist bound structures as compared to the antagonist bound structures (Figure 3). As agonist binding leads to the activation of CB_1_, active and inactive structures have distinct N-terminus feature values. The exception to this rule is the agonist and Negative Allosteric Modulator (NAM) bound structure (PDB ID: 6KQI^14^), where the structure remains inactive because of NAM binding. On the other hand, in the active and inactive structure, N-terminus always remain above the orthosteric pocket; therefore, N-terminus feature values do not distinguish between active and inactive crystal structures for CB_2_ (Figure 2 and 3).
2. In CB_1_, due to the extended structure of the antagonist, TM1 moves outwards towards the membrane in the inactive structures (Figure 2), whereas extracellular TM1 conformation of CB_2_ stays relatively similar in the active and inactive state in the protein. The higher variation in extracellular TM1 movement between active and inactive structures in CB_1_ as compared to CB_2_ is shown in Figure 3.
3. Another major difference in activation of CB_1_ and CB_2_ is observed in toggle switch residue movement. For CB_1_, twin toggle switch residues (W^6.48^ and F^3.36^) undergo translational changes (Figure 2). During the activation, W^6.48^ moves towards the TM5, which leads to the displacement of F^3.36^ towards TM2. The displacement is captured by the difference in the center of mass of the sidechain of these two residues in the z-direction (Figure S2). In the case of CB_2_, F^3.36^ does not undergo conformational change. Instead, agonist binding to CB_2_ has shown to rotate the toggle switch residue W^6.48^.^16^ This rotational motion is captured by calculating the *χ*_2_ angle of W^6.48^. Projection of translational and rotational movement features of toggle switch show that former feature could capture the distinction between active and inactive structures for CB_1_ and latter for CB_2_ (Figure 3).
4. For CB_2_, inactive structures in intracellular TM5, the sidechain of Y^5.58^ faces towards the membrane and creates more space for TM6 to move inside in the inactive structure. In the active structure, Y^5.58^ side chain (I^2.43^-Y^5.58^ distance) moves inside the transmembrane domain and faces towards the TM2 (Figure 2). No such conformational change is observed for CB_1_. Y^5.58^ always remains inside the transmembrane domain.

**Table 1:** Comparisons of conformational feature values between apo (without ligand) and holo (with agonist and antagonist) simulations of CB_1_ using mean difference and K-L divergence.

| Features | $\vert E(Holo_{inactive}) - E(apo)\vert + \vert E(Holo_{active}) - E(apo)\vert $ | K-L divergence (Holo<sub>active</sub> and apo) | K-L divergence (Holo<sub>inactive</sub> and apo) |
| --- | --- | --- | --- |
| N-terminus Movement | 0.4 | 2.1 | 1.3 |
| Extracellular TM1 movement | 0.2 | 1.9 | 1.1 |
| Toggle Switch translational movement | 0.3 | 1.3 | 1.8 |
| Toggle Switch rotational movement | 0.1 | 0.3 | 0.4 |
| Intracellular TM5 movement | 0.0 | 0.7 | 0.9 |
| Intracellular TM6 movement | 0.3 | 1.4 | 2.2 |
| Intracellular TM7 movement | 0.4 | 0.3 | 1.7 |

**Table 2:** Comparisons of conformational feature values between apo (without ligand) and holo (with agonist and antagonist) simulations of CB_2_ using mean difference and K-L divergence.

| Features | $\vert E(Holo_{inactive}) - E(apo)\vert + \vert E(Holo_{active}) - E(apo)\vert $ | K-L divergence (Holo<sub>active</sub> and apo) | K-L divergence (Holo<sub>inactive</sub> and apo) |
| --- | --- | --- | --- |
| N-terminus Movement | 0.1 | 2.1 | 0.7 |
| Extracellular TM1 movement | 0.3 | 0.9 | 1.1 |
| Toggle Switch translational movement | 0.1 | 1.5 | 1.7 |
| Toggle Switch rotational movement | 0.2 | 0.8 | 2.4 |
| Intracellular TM5 movement | 0.5 | 0.4 | 1.5 |
| Intracellular TM6 movement | 0.2 | 1.0 | 1.7 |
| Intracellular TM7 movement | 0.5 | 0.1 | 2.9 |

### Thermodynamic relevance of structural features in cannabinoid receptors’ activation

All extracellular (N-terminus and TM1 movements), transmembrane (Toggle Switch translation and rotation), and intracellular features (TM5, TM6 and TM7 movements) that distinguish active and inactive states of the receptors identified from different crystal structures may not be thermodynamically and kinetically relevant (Figure 3). Experimentally determined side chain conformations occasionally do not have good resolutions. Hence, it may over or under emphasize some of the conformational changes. Furthurmore, features that do not distinguish between active and inactive state for respective proteins (e.g., Toggle switch rotation for CB_1_; TM1 movement for CB_2_) may obtain distinct stabilized values in intermediate states during activation. Therefore, to check the thermodynamic relevance of each feature in CB_1_ and CB_2_ conformational ensemble, we compared the apo (without ligand bound) and holo (with ligand bound) simulation of both proteins.

Compairing ∼ 20*μ*s of unbiased MD simulation for agonist bound active and antagonist bound inactive states of both CB_1_ and CB_2_ (Table S2), it is observed that the conformational ensemble of every feature remains stable (Figure S3). It reinforces that the features observed in the crystal structures are thermodynamically relevant. The features that distinguish between CB_1_ and CB_2_ activation in experimentally determined structures are stabilized with different feature values.

The presence of intermediate states was judged by comparing the conformational ensemble of important features obtained from apo simulations with the ligand-bound active and inactive holo simulation (Figure S3). Simulation details are shown in Table S2. Three metrices are used to judge the feature importance : (1) sum of the absolute value of the mean difference between apo and holo simulations | *E*(*Holo*_*inactive*_) − *E*(*apo*) | + | *E*(*Holo*_*active*_) − *E*(*apo*) |, where E(Feature) represents the mean of the feature from simulation. (2) KullbackLeibler divergence (K-L divergence) of apo simulation with respect to agonist bound active simulation (3) K-L divergence of apo simulations with respect to agonist bound active simulations.^47^ Here, features were normalized for better comparison using these metrices. As expected, all the features which show distinction in crystal structures for both proteins have larger values as shown in Table 1 and 2. This indicates that these features undergo large changes during activation. On the other hand, features that were found to be only important for CB_1_ or CB_2_ show higher values for the metrices for only the respective protein. The only exception to this rule is the extracellular TM1 movement for CB_2_, where ligand-bound active and inactive conformational ensembles show overlapping distribution, whereas apo activation conformational changes much wider distribution of feature values (Figure S3), which may reveal the newer conformations. Therefore, using our simulation, we observe changes in the conformational features of the proteins that are not deemed important from static crystal structures. Overall, for CB_1_, features that undergoes conformational changes during activations are N-terminus distance, extracellular TM1 movement, toggle switch translational movement, intracellular TM6 and TM7 movements. Contrastingly for CB_2_, conformational changes are observed in extracellular TM1 movement, toggle switch rotational movement, intracellular TM5, TM6 and TM7 movements.

### Allosteric communications between CB_1_ and CB_2_ structural features

In the active state, conformational changes in the intracellular part of GPCR proteins help to transduce signals by secondary messenger (G protein and *β*-arrestin).^20,48^ These intracellular changes are allosterically linked with extracellular and transmembrane regions of protein. ^19^ To observe how the conserved and non-conserved changes in CB_1_ and CB_2_ are allosterically connected, we predicted whether the change in one feature leads to the change in the other features. To estimate this, the absolute difference of *P* (*Feature*_*active*_|*Condition*_*active*_) and *P* (*Feature*_*active*_|*Condition*_*inactive*_) was determined for every combination of features. These conditional probabilities were measured by discretizing the features based on a threshold that divides the features into active and inactive. The threshold values are taken as an average of active and inactive crystal structures. The obtained probabilities were weighed based on the MSM estimated equilibrium probabilities of each state.

Based on the calculations for CB_1_, it shows that N-terminus movement is highly correlated with all the extracellular (TM1 movement), transmembrane. (Toggle switch translational movement), and intracellular features (TM6 and TM7 movement), showing the importance of the N-terminus movement in activation (Figure 4A and Figure S4A). Projection of MSM weighted free energy landscapes show that with the N-terminus inside the pocket, other features remain mostly in the stabilized inactive state (Figure S5A, S5B, S5C, and S5D). On the other hand, extracellular TM1 movement is not strongly coupled with transmembrane and intracellular features (Figure 4A and S4B). Free energy projection of TM1 movement with respect to intracellular and transmembrane features shows those features are able to obtain active and inactive forms without depending on the TM1 value (Figure S5E, S5F, S5G). Similarly, toggle switch translational movement also shows relatively smaller values of coupling for intracellular TM6 movements and TM7 movements (Figure 4A and S4C). However, the MSM-weighted free energy landscape can explain this relatively smaller value. The free energy plots show an L-shaped landscape between the toggle switch and intracellular features depicting the change to be sequential (Figure S5H and S5I). Thus, changes in the toggle switch can lead to changes in the intracellular features. Lastly, both conditional probability difference and free energy landscape show the intracellular features are highly coupled with each other, which depicts that the TM6 movement will lead to the TM7 movement (Figure 4A, S4D, S4E, and S5J).

**Figure 4:**
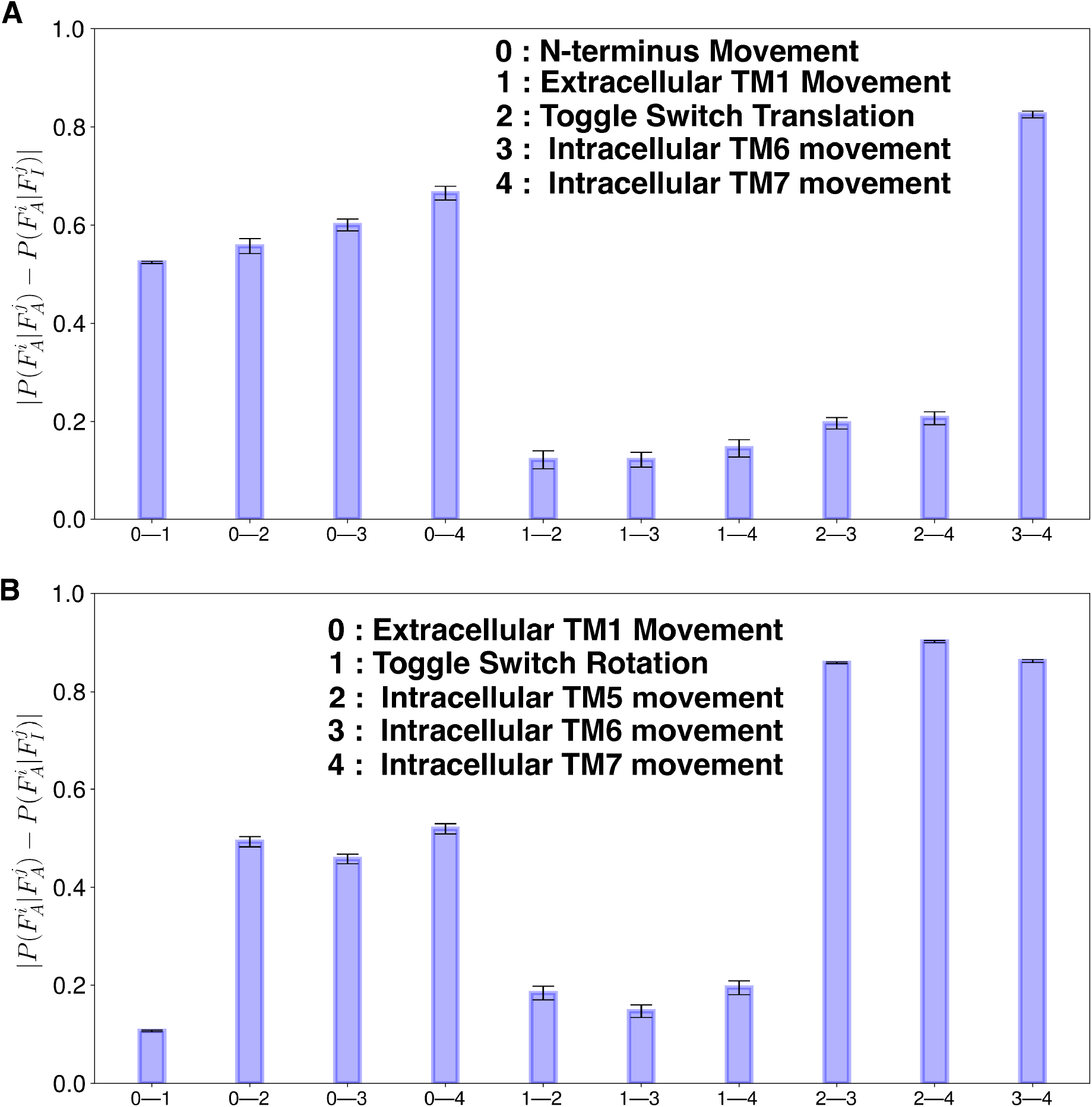
Absolute differences of conditional probabilities 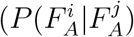 and 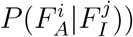 of structural features for CB_1_ (A) and CB_2_ (B). 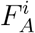 denotes the *i*th feature is in active state. 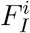 denotes the *i*th feature is in inactive state. Errors in the conditional probability calculations were calculated based on bootstrapping. 20 bootstrap samples were selected with 80% of the total number of trajectories.

For the CB_2_, all the intracellular features are highly coupled with the extracellular TM1 movement (Figure 4B and S6A). MSM weighted free energy landscape shows that when intracellular features remains in the inactive states, extracellular TM1 can have a wider range of movement. Whereas, in active state, extracellular TM1 movement is restricted (Figure S7A, S7B, S7C, S7D). From the experimentally obtained structures, it has been shown that toggle switch residue rotates when the agonist molecule binds to the receptor. Therefore, it has been hypothesized that toggle switch rotation caused by agonists leads to intracellular changes in the CB_2_ receptor.^15,16^ However, our calculations show that toggle switch rotation has minimal effect on the intracellular movement (Figure 4B and S6B). From the Free energy projection, it is also clear that intracellular features can adopt both active and inactive forms irrespective of toggle switch rotation feature value (Figure S7E, S7F, S7G). Agonist-bound CB_2_ structure (PDB ID: 6KPC) remains in an inactive state as there is no intracellular movement observed, which supports our finding that there is a lack of correlation between toggle switch rotation and intracellular movement.^16^ For CB_2_, intracellular TM5 movement is shown to be important for canonical GPCR activation. The free energy landscape shows that L shaped landscape between intracellular TM5 movement with respect to the intracellular TM6 and TM7 movement, which depicts that TM5 movements may be the cause of the activation of the GPCR, not the toggle switch rotation (Figure S7H and S7I). Lastly, similar to CB_1_, intracellular TM6 and TM7 movement are highly correlated with each other (Figure 4B and S7J). Therefore, these analyses show that most of these features (Except for TM1 movement for CB_1_; toggle switch rotation in CB_2_), which are physically far away in protein topology, are allosterically influence conformational changes of other features resulting in protein activation.

### Activation mechanism for CB_1_ and CB_2_

Previous sections show the important conformational changes for CB_1_ and CB_2_ during the activation. However, individual feature movements or projections of a few features do not reveal the activation mechanism of the receptors and intermediate states involved in the process. To better understand the activation mechanism, we clustered the protein conformational ensemble using neural network architecture, VAMPnets.^46^ VAMPnets architecture discretizes the protein conformational space into metastable states by building a coarse-grain Markov state model (MSM). The number of metastable states is decided by minimizing the error in implied timescale with a condition that each metastable state will have minimum population of 4% of entire ensemble (Figure S8, S9, S10, S11). Using these criteria, we built a six-state model for both CB_1_ and CB_2_. These six states are projected on the tic1 and tic2 dimensions, which are linear combinations of the slowest features (Figure 5 and 6). Here, two of the metastable states are presented as active and inactive based on the projection of the active and inactive experimentally determined structures onto tics. Except for the active and inactive states, there are four intermediate metastable states obtained from the model, named I1, I2, I3, and I4 for both CB_1_ and CB_2_ (Figure 5 and 6). For CB_1_, I1 and I2 are structurally similar to the inactive state of the receptor (Table S3 and Figure S12). Whereas, I3 and I4 are structurally similar to active state (Table S3 and Figure S12). The structural differences between different metastable states are calculated using K-L divergence based on the closest heavy atom distances between all residue pairs with respect to the inactive state. In Figure 5 and 6, the calculated K-L divergence is shown as a color bar on the representative protein structure, where the red represents structural divergence. Furthermore, normalized values of the important features (that have been discussed in the earlier section) are calculated for each metastable state as shown in Figure 5 and 6. Transition kinetics between the metastate states are shown as mean free passage time (MFPT) if there is a direct transition. Using the above analyses, it is observed that from the CB_1_ inactive state, both I1 and I2 transitions are possible. I1 transitions are kinetically more favorable compared to I2. Inactive to I1 transition involves the movement of intracellular TM1 region as shown in K-L divergence analysis and bar plot (Figure 5). In the case of inactive to I2 and I1 to I2, the transition involves the movement of the N-terminus in the upward direction. Movement of the N-terminus is a slow timescale process as these movements are related to the first eigenvectors of MSM (Figure S13). Intracellular and transmembrane regions of I1 and I2 metastable states remain similar to inactive states. Conformations of the I2 metastable state are similar to the agonist, and NAM-bound crystal structure (PDB ID: 6KQI^14^), where the structure remains in the inactive state in spite of the binding of the agonist in the pocket (Figure S14). I1 to I3 transition involves rearrangement of the intracellular and transmembrane features as N-terminus starts to move out of the pocket. K-L divergence analysis and bar plot show that the major difference occurs in N-terminus, toggle switch region, intracellular TM6, and TM7 region (Figure 5 and S12). This step is the the kinetically slowest process during the activation. Once the intracellular and transmembrane rearrangement happens in I3, the rearrangement of structural rearrangement happens between I3 & I4 and I3 & active state of the receptor. In the case of the I3 to I4 transition, N-terminus moves slightly further upward the pocket, and further intracellular features movement happens (Figure 5). During I3 to active state transition, the N-terminus moves out of the binding pocket, which TM1 to come inside the pocket; however, intracellular features remain relatively the same (Figure 5). Between these two transitions, the later transition is kinetically faster as I3 to I4 transition involves N-terminus movement (Figure 5).

**Figure 5:**
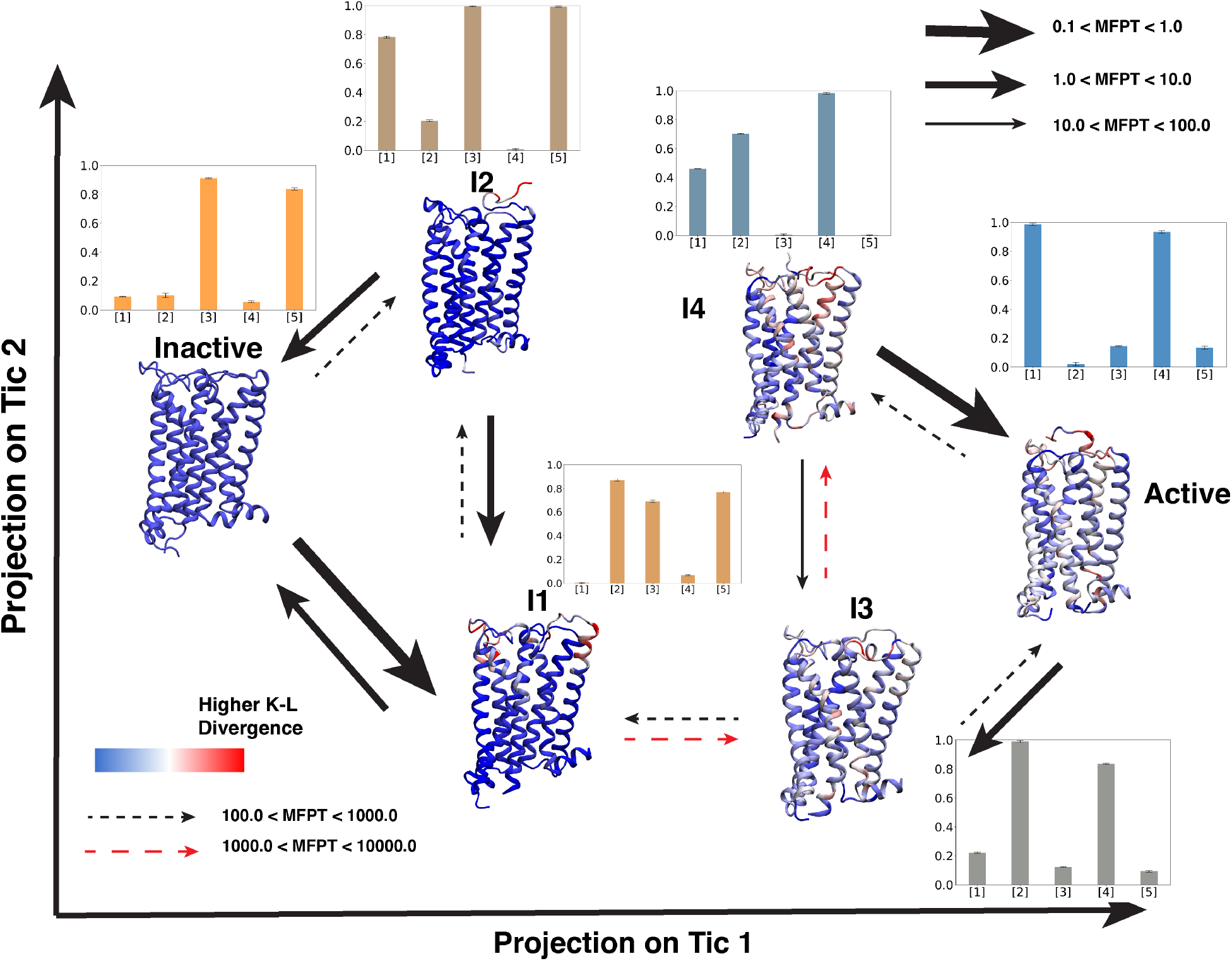
Representative structures from six CB_1_ metastable states obtained from VAMPnets are projected on tic 1 and tic 2 spaces. Cartoon representations of receptors are colored based on K-L divergence. K-L divergence of each metastable state was calculated compared to inactive metastable state. Transition between metastable states are shown as arrows. Thickness of the arrows represents timescale of the transition which was calculated using transition path theory (TPT). Bar plots per metastable state shows the normalized values of important structural features, where [1], [2], [3], [4], [5] represent N-terminus movement, extracellular TM1 movement, toggle switch translational movement, intracellular TM6 movement, intracellular TM7 movement respectively. Errors in bar plots were calculated based on bootstrapping. 20 bootstrap samples were prepared by selecting 1000 frames from each metastable states.

**Figure 6:**
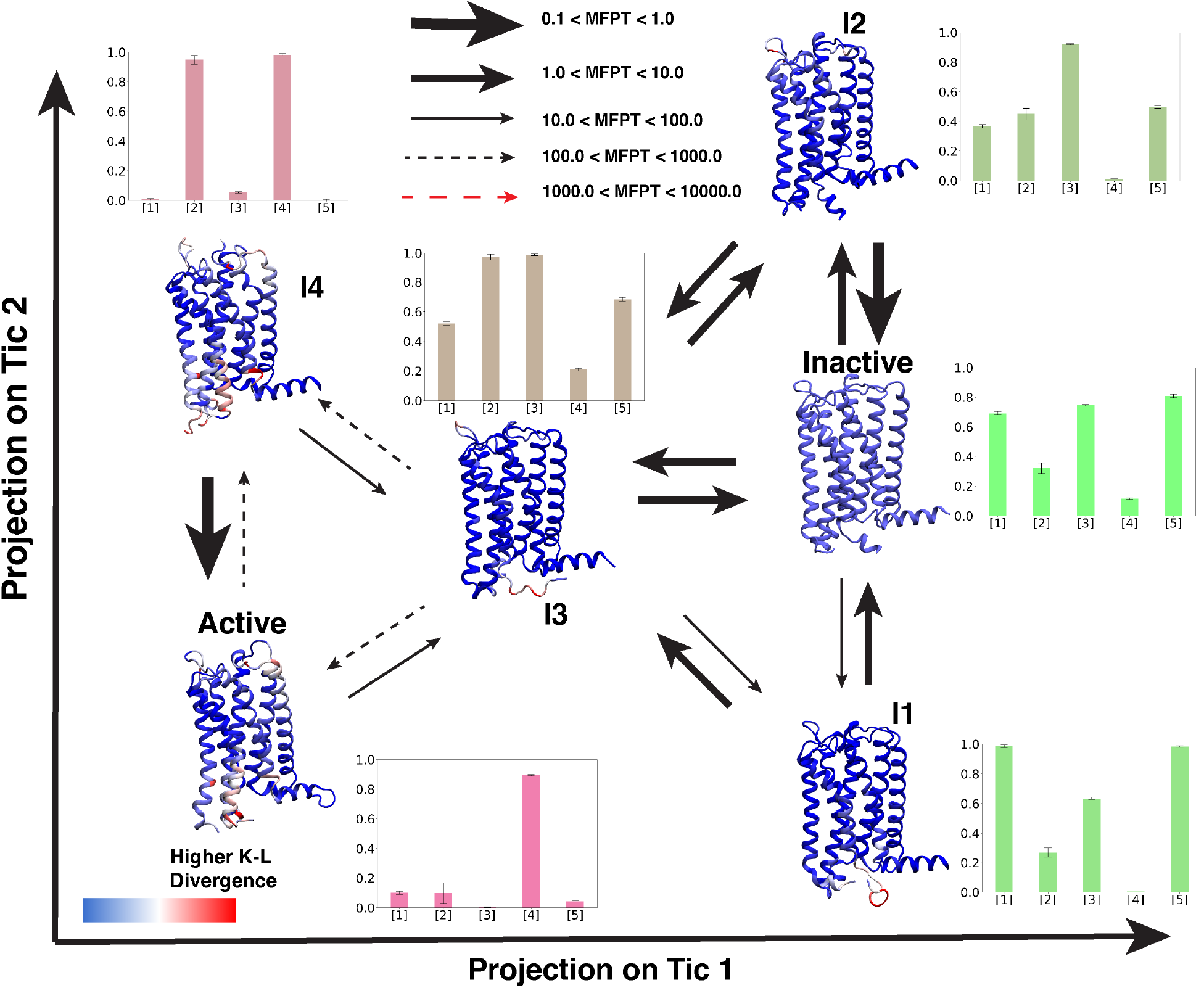
Representative structures from six CB_2_ metastable states obtained from VAMPnets are projected on tic 1 and tic 2 spaces. Cartoon representations of receptors are colored based on K-L divergence. K-L divergence of each metastable state was calculated compared to inactive metastable state. Transition between metastable states are shown as arrows. Thickness of the arrows represents timescale of the transition which was calculated using transition path theory (TPT). Bar plots per metastable state shows the normalized values of important structural features, where [1], [2], [3], [4], [5] represent extracellular TM1 movement, toggle switch rotational movement, intracellular TM5 movement, intracellular TM6 movement, intracellular TM7 movement respectively. Errors in bar plots were calculated based on bootstrapping. 20 bootstrap samples were prepared by selecting 1000 frames from each metastable states.

For CB_2_, intermediate states I1, I2, and I3 are structurally and kinetically similar to the inactive state of the receptor, whereas I4 is structurally similar to the active state (Table S4; Figure 6 and S15). K-L divergence shows no major structural changes between the intermediate I1 and I2 & inactive metastable states. Therefore, these states can interconvert with a faster timescale between each other (Figure 6). The inactive and I3 states are separated in tic one space. Intracellular TM6 and TM7 features are correlated in the tic 1 space (Figure S16). In this transition, TM6 moves out slightly, whereas Y^7.53^ moves towards the TM5 (Table S4 and Figure S15). From I3 to I4 and active state movement, this TM6 movement leads the Y^5.58^ to move inside the transmembrane region. Both of these transitions are kinetically slower as it involves rearrangement of the intracellular regions of the CB_2_.

MFPT calculations based on transition path theory (TPT) clearly show that the transition between inactive to active state in CB_2_ is much faster than CB_1_. A Kinetic monte Carlo (kMC) study on MSM based transition matrices also verifies that inactive to active transition is faster for CB_2_ transition (Figure S17A and S17B). kMC analysis also indicates that transition between inactive, I1, and I2 is kinetically much faster for CB_1_, compared to I1 to I3 transition. I1 to I3 transition leads to the protein moving from the inactive to the active region. Similarly, the kMC study verifies that inactive, I1, I2, and I3 are kinetically closer to each other for CB_2_. Therefore, the transitions between the states are relatively faster. Once the transition happens from I3 to active or active like I4, it remains stabilized in active-like proteins. The above analyses reveal intermediate states during CB_1_ and CB_2_ activation, by which the sequential conformational change happens that leads the protein from the inactive to the active state, and the distinction between changes between CB_1_ and CB_2_.

We further compared how the conformational changes in the distinct metastable states lead to different allosteric communication networks. Allosteric pipelines in different metastable states were obtained following the analyses described in Bhattacharya and Vaidehi.^49^ From these analyses, major distinctions in the allosteric pipelines were observed for CB_1_ in the inactive and inactive intermediate states (I1 and I2) compared to active and active intermediate states (I3 and I4) (Figure S20). In inactive, I1, and I2 states, extracellular TM1, and TM2 are allosterically connected to ECL2 via N-terminus. The downward movement of the N-terminus can be explained to be responsible for these allosteric pipelines as there is more interaction between N-terminus and ECL2. There are fewer pipelines connecting extracellular and intracellular domains. In contrast, major pipelines are observed connecting these two domains in active and active intermediate states. Especially all these three states have an allosteric pipeline through TM6, which indicates the outward movement of TM6 helps to create an allosteric path. On the other hand, for CB_2_, no such clear distinction is observed in allosteric pipelines between active like (active, I4) and inactive like states (Inactive, I1, I2, and I3). Most of the metastable states contains extracellular to intracellular communications. This contrasting network pattern in CB_2_ compared to CB_1_ can be explained by the lack of the major extracellular conformational changes in CB_2_. Hence, these analyses show how conformational changes and the consequent change in allosteric communications facilitate receptor activation.

### Explaining selectivity of Cannabinoid receptors

Ligand binding to a receptor can be explained using two distinct mechanisms, i.e., conformational selection and induced fit.^50^ In the induced fit mechanism, ligand binding leads to conformational changes in the receptor. On the other hand, the conformational selection mechanism sees the ligand preferentially bind to a particular protein conformation. As active and inactive binding pocket volumes of experimentally obtained CB_2_ structures are relatively similar, contrasting to the dissimilarity between CB_1_ structures, we hypothesize that the CB_2_ pocket shape remains similar throughout the activation process. If so, then the binding of CB_2_ selective agonists would demonstrate a conformational selection mechanism in preference to this generalized binding pocket conformation seen during CB_2_ activation. On the contrary, the CB_1_ binding pocket shape changes with the movement of the N-terminus during the activation. Therefore, these selective ligands can only bind to certain metastable states for CB_1_ which might have similar pocket characteristics as of CB_2_, decreasing the overall affinity. To test this hypothesis, we first calculated the binding pocket volumes of the metastable states during CB_1_ and CB_2_ activation. Our analyses show that CB_1_ metastable states have different binding pocket volumes based on the position of the N-terminus (Figure 7A). As the N-terminus remains completely inside the pocket during inactive and I1 states, agonist binding volume is comparatively smaller compared to the other four metastable states. Thus, as the CB_1_ conformations differ, an induced fit mechanism must follow so that CB_1_ can appropriately bind to a CB_2_ selective ligand. In the case of CB_2_, the binding pocket volume does not change during the activation due to a lack of N-terminus motion (Figure 7B). Here our results illustrate how CB_2_ maintains a competent conformational state for selective ligand binding throughout all stages of activation.

**Figure 7:**
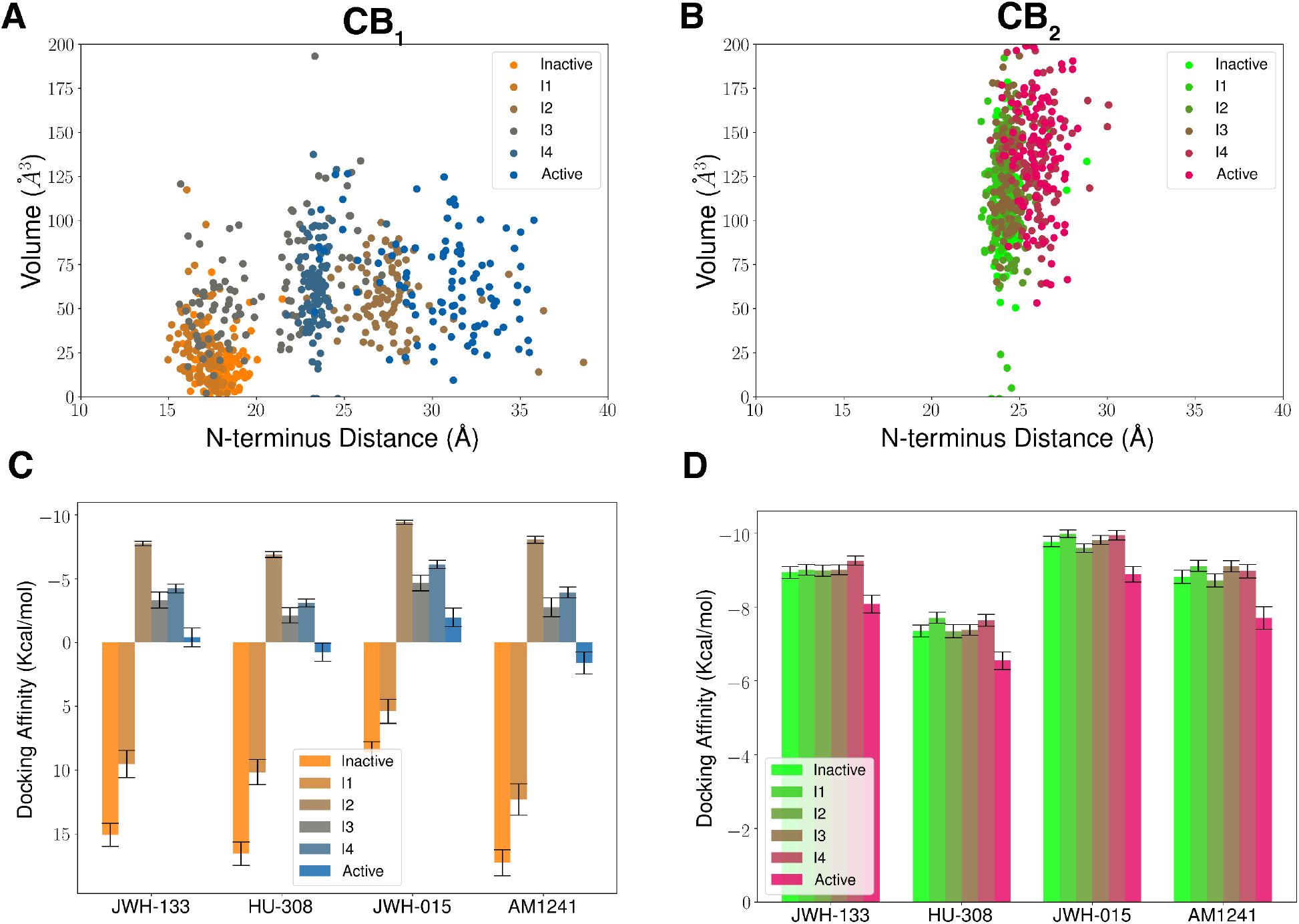
Binding pocket volumes for each metastable state of CB_1_ (A) and CB_2_ (B) are plotted against N-terminus distance as a scatter plot. Docking affnity (kcal/mol) of the four CB_2_ selective agonists for each metastable state of CB_1_ (A) and CB_2_ (B) are plotted as a bar plot. Docking and volume calculations were performed on 100 structures from each metastable state selected based on the MSM probabilities. For each protein structure, docking affnity is obtained as an average of top three dock poses. Error shown in docking calculations is the standard error which the standard deviation of docking affnity distribution obtained from 100 structures divided by 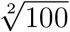.

To find whether CB_2_ selective agonist binding affinity depends on the binding pocket shape, we docked CB_2_ selective ligands into CB_1_ and CB_2_ metastable states (described in the method sections). Four CB_2_ selective ligands (JWH-133, HU-308, JWH-015, AM-1241) are selected from the literature,^32^ which shows at least ten-fold selectivity (Figure S18). Docking results of CB_2_ selective ligands to CB_1_ show that docking affinity differs for different metastable states (Figure 7C). For inactive and I1 states where the binding pocket volume is small, the docking is very hard due to the steric hindrance of the N-terminus. In the other four states, docking results show variational docking affinity, with I2 states having the highest affinity for all the CB_2_ selective ligands compared to the active state. Therefore, in the CB_1_ active state, these ligands have a less binding probability. This differential binding affinity is explained by the differential binding pose of the ligands in I2 and active states. In I2, there is a larger gap between TM2 and ECL2, which allow the ligands to bind between the space of ECL2, TM1, and TM2 (Figures S19A and S19B). In the active state, the ligands bind deep in the pocket due to the movement of the toggle switch. Toggle switch movement creates a space for ligand to bind (Figures 8A, 8B, 8C, and 8D). Whereas the docking affinities for these ligands are similar to each metastable state for CB_2_ as the receptor pocket shape remains relatively similar (Figure 7D). Therefore, these analyses show that how the difference in pocket shape due to distinct activation mechanism of CB_1_ and CB_2_ may lead to ligand selectivity towards CB_2_.

**Figure 8:**
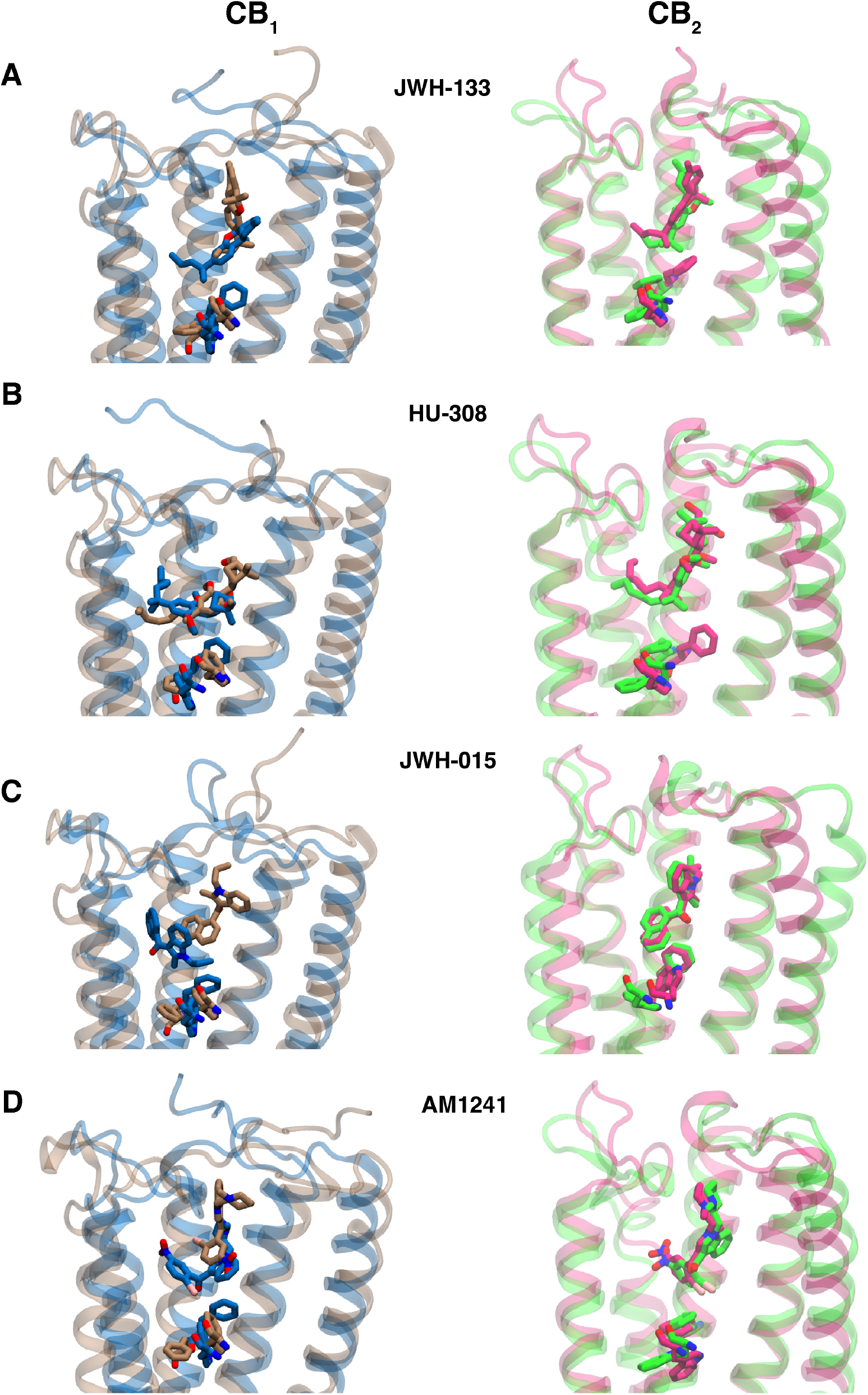
Differences in most stable docking poses between active and I2 metastable states for CB_1_ (Left panel) for four different CB_2_ selective ligands. Differences in most stable docking poses between active and inactive metastable states for CB_2_ (Left panel) for four different CB_2_ selective ligands.

## Conclusions

CB_2_ selective agonists are emerging as potential drug targets for treating inflammation. A number of CB_2_ selective agonists have entered clinical trial phase.^34,35^ However, none of these molecules has been approved as a drug by FDA. There are two of the major drawbacks in developing CB_2_ selective drugs. First, high lipophilic nature of the CB_2_ selective agonists are not suitable to be drug target. Second, some CB_1_ activity may retain in those molecules which leads to off-target side-effect. In this work, we mechanistically explain the selectivity of CB_2_ selective agonists by hypothesizing that reason for selectivity lies in the difference of activation mechanism. Here, we have studied the activation mechanism of both CB_1_ and CB_2_ by with combinations of using molecular dynamics, Markov state model and neural network.

Our results showed three major distinctions in conformational changes in both receptors. (1) In the extracellular region due to the movement of the N-terminus, we observe a large change in the binding pocket volume during the activation of the CB_1_, whereas, for CB_2_ there are hardly any changes in the binding pocket. (2) For CB_1_, toggle switch W^6.48^ undergoes translational changes during the activation mechanism, which lead to the changes in the intracellular matrices. While, for CB_2_, toggle switch W^6.48^ only rotates in the same position. Our analysis also shows that toggle switch rotation does not effect the movement intracellular movements for CB_2_. (3) Intracellular TM5 movement of CB_2_ affects the TM6 and TM7 movement critical for receptor activation. On the other hand, no major conformational change is observed in intracellular TM5 of CB_1_.

Vampnets analysis further reveals the metastable states and sequential transition of the structural features during the activation. During the CB_1_ activation, initial extracellular movements of N-terminus, TM1 lead to the change in further change in the transmembrane and intracellular features. Conversely, due to lack of the extracellular changes, the activation of CB_2_ is kinetically faster compared to the CB_1_.

Further analyses show that due to the lack of the extracellular movement during CB_2_ activation, binding pocket volume remains relatively same for CB_2_. Docking of metastable states show that CB_2_ selective agonists show the similar high binding affinities for each metastable states of CB_2_, depicting the preferential binding of these ligands in the generalized CB_2_ binding conformation. Whereas, these ligands can only be selectively bind to few metastable states in CB_1_ due to the larger conformational change in the CB_1_ binding pocket. This explains lack of overall binding affinity of these ligands towards CB_1_. Overall, we provide mechanistic knowledge about cannabinoid receptor selectivity by studying the activation mechanism. This study will guide to design new CB_2_ selective agonists with better druggability profile by acting as virtual screening criteria for these ligands.

## Methods

### System Preparation

To perform apo (without ligand) simulation, we started molecular dynamics simulations from inactive (CB_1_ inactive state PDBID: 5TGZ;^9^ CB_2_ inactive state PDBID: 5ZTY^12^) and active (CB_1_ active state PDBID: 5XRA;^11^ CB_2_ active state PDBID: 6KPF^16^) state structures of CB_1_ and CB_2_. Non-protein residues were deleted from the CB_1_ and CB_2_ crystal structure. The thermostabilized mutations were mutated back to original residue using tleap. Hydrogen atoms were added to protein amino acid residues using reduce command of AMBER package.^51^ To add consistency in the residue numbering, 4 residues (CB_1_ residue number 99-103) were added in the truncated N-terminus for CB_1_ active structure. Intracellular region between TM5 and TM6 was truncated for both CB_1_ and CB_2_. Terminal end of N-terminus and C-terminus & unconnected residues of TM5 and TM6 were neutralized with ACE and NME residue. Both CB_1_ and CB_2_ was embedded in POPC bilayer with 150mM NaCl solution in both extracellular and intracellular direction. TIP3P water model was used for MD system.^52^ AMBER ff14SB and lipid17 forcefield was used for protein and lipid parameterization.^53,54^ Agonist bound active (CB_1_ agonist: AM11542, CB_2_ agonist: AM841) and antagonist inactive (CB_1_ antagonist: AM6538, CB_2_ antagonist: AM10257) holo simulation were also performed. GAFF forcefield parameters for the ligands are obtained using antechamber.^55,56^ Similar modifications were made in the proteins using tleap and embedded in bilayer using CHARMM-GUI.^57^

### Simulation Details

Atomistic unbiased molecular dynamics simulation was performed with AMBER MD engine.^58^ Before running the production MD run, prepared MD systems were minimized and equilibrated. Minimization was performed for 15000 steps; where the systems were minimized for 5000 steps with gradient descent algorithm and followed by conjugate gradient algorithm for the rest 10000 steps. Minimized systems was heated and pressurized sequentially in NVT and NPT ensemble, respectively. Heating step was performed for 2 ns where system was heated from 0K to 300K in NVT ensemble. Anaisotropic pressure was applied to the heated systems to reach the constant pressure of 1 bar with NPT ensemble. Pressurization step was also performed for 2 ns. Berendsen thermostat and barostat was used to fix the temperature and pressure.^59^ During the heating and pressurization, protein backbone for each system was restrainted using a spring force with a spring constant of 10 kcal/mol/ Å ^2^. Subsequently, restraint was removed and each system was equilibrated for 46 ns for collecting data. Numerical integration was performed using verlet algorithm to update velocity and coordinate. Time step for each numerical integration is 2 fs. Shake algorithm was implemented by constraint bond movement of the hydrogens to bonded to heavy atoms.^60^ Periodic boundary condition was applied to each MD system. Particle mesh Ewald method (PME) was implemented for long-range electrostatic calculation.^61^

### Featurization of Protein Conformational Ensemble

To perform adaptive sampling and markov state model (MSM) building (Discussed in the following sections), protein conformations in MD system need to be represented by the descriptors that can capture protein conformational changes. To find the descriptors, we have implemented residue-residue contact score (RRCS) analysis.^62^ Using RRCS method, a score was assigned to each residue pair distances for a particular protein conformation which are more than one helical tern away. This score was calculated based on the every atom pair distances between two residues following the same algorithm of Zhou et al. ^62^ To find the major structural changes during activation for CB_1_ and CB_2_, we compared the change in RRCS between the active and inactive structure. *α*-carbon distances of the residue pairs, where the RRCS is changing more than 3, were used as a feature to describe a protein conformation (Table S5 and S6). As shown in Figure S22, these descriptors can capture the extracellular, transmembrane and intracellular changes for both CB_1_ and CB_2_.

### Adaptive Sampling

Previous experimental and computational studies depict that GPCR activation occurs in a microsecond to millisecond timescale.^20,63–65^ However, each numerical integration time step for MD simulation is of femtosecond timescale.^66^ Therefore, capturing the entire activation process by running a single long trajectory is time consuming process. Furthermore, to capture the thermodynamic and kinetics of the conformational changes better sampling is needed in high energy transition states.^67^ Thus, to capture the entire protein conformational ensemble, adaptive sampling protocol was implemented.^67–70^ Adaptive sampling is a well established sampling technique used for studying ligand binding, ^71–73^ protein conformational change^63,74–76^ and ligand selectivity.^77^ First, conformations obtained from MD simulations are expressed in terms of collective variables. Second, collective variable space are clustered into different states using k-means clustering. Third, based on the population of the each state, MD frames are selected from least populated clusters to start simulation for the next round. GPCRs consist of the multiple allosterically coupled conversed motifs distributed in extracellular, transmembrane and intracellular regions. These motifs structurally rearrange themselves when protein conformational change happens from inactive to active. Therefore, finding suitable adaptive sampling matrices to minimize the amount of simulation time is difficult. Here, we used our preexisting knowledge of the structural changes to select CVs and modify the CV based on the requirement. For initial round of sampling, we selected the extracellular TM1-TM6 distance and intracellular TM3-TM6 distance as a collective variable for adaptive sampling. For both CB_1_ and CB_2_, sampling was performed from both active and inactive structures until energy landscape was connected. Next, 24 and 20 distinct features (Extracellular distances, Toggle switch movements, Intracellular TM6 and TM7 movements) were selected for CB_1_ and CB_2_ to improve the speed of the sampling (Table S7 and S8). To check whether simulations able to observe the transition between the inactive to active states in the RRCS descriptor space (as discussed in previous section), we performed dimensionality reduction using time independent component analysis. On the projection of slowest two tic components, transition between active and inactive state was observed in CB_2_ conformational ensemble. However, there was a disconnect between active and inactive minima in tic projection of CB_1_. Therefore, additional rounds of sampling were performed from the clusters that are closest from each other in the two disconnected regions to observe the transition between active and inactive minima is observed for CB_1_. In total, 490 and 278 *μ*s simulations were performed to capture apo activation process for CB_1_ and CB_2_. Adaptive sampling for holo systems is performed on the 24, 20 structurally known features for CB_1_ and CB_2_ holo systems for approximately 20 *μ*s.

### Markov State Models

From adaptive sampling, we generate plethora of short trajectories that only capture the small changes in protein conformation. Markov state model was developed to connect the information from individual trajectories to by building transition network between distinct states.^78–80^ In markovian processes, the future step of the process should only depend on present states, not on the past states.^81^ Molecular dynamic simulations follow the same principle as the position and velocity of the future step is calculated from the energy and force calculation based on the current state. From MSM, we obtain stationary probability distribution of the protein ensemble and timescales of the slowest process by calculating transition probability matrix (TM). TM calculates probability of transition between discretized protein space by calculating the jump between different states after lag time. Lag time is the minimum time at which markovian property of the discretized states is valid. Building markov state model consist of four different steps: featurization, dimensionality reduction, clustering and hyperparameter optimization. Featurization of the protein conformational space was done from the distances calculated from RRCS (discussed in previous subsection). Dimensionality reduction was performed on the featurized space with TICA analysis. MSM captures the slowest timescale processes. Using TICA, we obtain new representations of features which is a linear combination of slowest features. Therefore, in the next discretization step, clustering algorithm can discretize the protein space which are distinct in slowest timescale. K-Means clustering algorithm was implemented in this case. The hyperparameters that needs to be optimized are lagtime at which markovian property of the discretized space is valid, number of tic dimensions on which clustering is performed and number of cluster components or microstates that can distinguish the slowest components. With a minimum lag time of 25 ns for both the systems, slowest timescales obtained from MSM converges, atleast in the log scale (Figures S23A and S23B). Hence, the markovian property is valid. To optimize other two hyperparameters, the VAMP-2 score was calculated.^82,83^ For reversible process, which follows the detailed balance, VAMP-2 score is the sum of the squares of the highest eigenvalues.^84^ MSM with the highest VAMP-2 score better captures the slowest timescale processes. The hyperparameters which maximizes the VAMP-2 scores are considered for the final MSM estimation (Figures S23C and S23D). Pyemma software package was used for entire MSM building process.^85^

### Trajectory Analysis

Features such as distances, angles are calculated using python library MDtraj.^86^ VMD software package was used for protein figure making and trajectory visualization.^87^ Ambertools CPPTraj was used for frame selection and trajectory modulations.^88^ All the analysis codes (e.g. conditional probability calculation) were written in python programming language and matplotlib library was used for plotting the figures as a graph. POVME software package was used for volume calculation of the binding pocket.^89^

### Metastable State Estimatation Using VAMPnets

Vampnets is a deep learning architecture used for building coarse grain MSM. ^46,84,90^ This architecture consists of two parallel deep learning networks which take the input feature values at time t and t+*τ*, where *τ* is the lagtime. This deep learning model is optimized by maximizing the VAMP-2 score. The output of the VAMPnet is the probability of the each frame belonging to a certain metastable state. VAMPnet was implemented using python library Deeptime.^91^ RRCS calculated features were as an input. Lagtime for the VAMPnet is selected as 25 ns. The number of the metastable states for each system was selected by the two creteria : (1) Each metastable state should have atleast 4 percent of the total population (Figure S10 and S11). (2) error bar in the timescale calculations is minimum (Figure S8 and S9). Based on these creteria, the number of metastable states chosen for CB_1_ and CB_2_ is six.

### Data-Driven Analysis

To capture the major structural changes between different macro states, closest heavy carbon atom distances was calculated between every residue pair of each frame. These calculations were performed on 1000 frames from each metastable state. These frames are selected based on the the probability of microstates belonging to the metastable states. Distance distribution between certain residue pair is compared to the identical residue pair distance distribution between distinct metastable states. Comparison is performed with symmetric K-L divergence analysis. To estimate the contribution of each residue, average K-L divergence of the all the residue pair distances was calculated that are associated with that residue.^47^

### Allosteric Pathway Calculation

To predict the allosteric pathway in different metastable states, similar procedure as explained in Bhattacharya and Vaidehi.^49^ First, normalized mutual information for each residue pair was calculated per metastable state based on dihedral angle movement. These calculations was performed on the 1000 frames as discussed in Data-driven analysis section. Based on the residue pairs, where closest heavy atom are within 5 Å, an undirected acyclic graph was created. Nodes in the graph denote a residue and edges are between the two residues which are within 5 Å of one another. Edge weight represents the MI_*max*_ MI_*ij*_, where MI_*ij*_ is the mutual information between the two nodes *i* and *j*. Shortest allosteric path was calculated for the residue pairs that are atleast 12 Å away from each other. Among all the calculated paths, 500 paths with highest mutual information were selected. These paths were clustered into optimal number of clusters based on the procedure explained in Bhattacharya and Vaidehi.^49^ Mutual information calculations have been performed with MDentropy software package.^92^

### Docking Calculations

To find the docking pose and the docking score of the CB_2_ selective ligands, in distinct metastable states of CB_1_ and CB_2_, four CB_2_ selective ligands were docked into binding pocket of distinct metastable states of CB_1_ and CB_2_. Four ligands were selected based on the criterion that those ligands show at least 10 fold more affinity for CB_2_ than CB_1_.^32^ Ligand structures were downloaded from pubchem using sdf format which is converted into the pdb format using Charmm-gui ligand builder.^57,93^ Amber tools Antechamber is used for converting the pdb format to mol2 format. ^55,56^ For docking, mol2 files were converted to pdbqt format. To represent each metastable state, 100 protein structures were selected from each metastable state proportional to the probability of microstates and converted to pdbqt format. Autodock-vina software was used for docking calculations.^94,95^ To specify the docking center, center of mass of alpha-carbon of binding pocket residues for CB_1_ and CB_2_ was selected Figure S24.

### Transition Path Theory

Transition path theory was implemented to calculate the effective timescale between metastable state transition.^96,97^ Effective timescale between two metastable states (A and B) is obtained from mean free passage time (1*/k*_*AB*_) as shown in equation 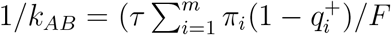, where F is the total flux between metastable state A to B and 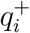 is the committer probability which is defined as the probability of microstate *i* reaching the metastable state B before A. A microstate is assigned to a metastable state based on the highest probability of belonging to each metastable state. Thus, six metastable states are defined as distinct microstates. TPT calculations were performed using Pyemma software. ^85^

### Error Calculation

To calculate errors for thermodynamic and kinetic quantities obtained from simulation, bootstrapping approach was followed. ^76,98^ For a certain calculation, N numbers of random bootstrap samples were generated with 80% of total trajectories. Based on the standard deviation of a calculated quantity from bootstrapped samples, error bars were generated. For example, for error calculation in free energy plots, MSM was created for 200 times using 80% of the original data to find the standard deviation in the probability density.

### Kinetic Monte Carlo Simulation

Based on the transition matrix of MSM, kinetic monte carlo (kMC) simulation was implemented to observe the evolution of features with time.^63,99^ kMC simulation is a stochastic process where in a given time step, next transition is selected based on the probability of the all possible transitions. The algorithm for KMC simulation is as follows: (1) Simulation starts at a particular microstate (obtained from clustering; as discussed in earliar section). (2) From the MSM transition matrix, the probabilities of all possible transitions (*P*_*i,j*_) from that microstate are obtained. As the sum of all transition probabilities is 1, a cumulative distribution (S) of possible transitions is created where 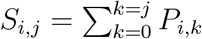. A random number (R) is generated between 0 to 1 to find the next possible transition. If R is between *S*_*i,j*_ and *S*_*i,j*+1_, *i*th state to *j* + 1th state transition is selected and simulation moves forward by lag time (*τ*). This algorithm is followed iteratively. In this case, kMC simulation was initialized from the inactive state of both the proteins.

## Acknowledgements

We thank Blue Waters sustained-petascale computing project, which is supported by the National Science Foundation (awards OCI-0725070 and ACI-1238993). D.S. acknowledges support from NIGMS MIRA award R35GM 142745. S.D. thank Balaji Selvam for help during the initial phase of the project. S.D. also thank Austin Weigle for suggestions during manuscript writing.

## Conflict of interest

The authors declare no conflict of interest.

## Code and data availability

All the necessary files (MSM object, bootstrap files, feature files, python scripts and protein conformations) that have been used to generate figures can be obtained from https://github.com/ShuklaGroup/Cannabinoid_activation.git.

## Notes

### Competing Interest Statement

The authors have declared no competing interest.

