## Supplementary Text for "Distinct Activation Mechanisms Regulate Subtype Selectivity of Cannabinoid Receptors"

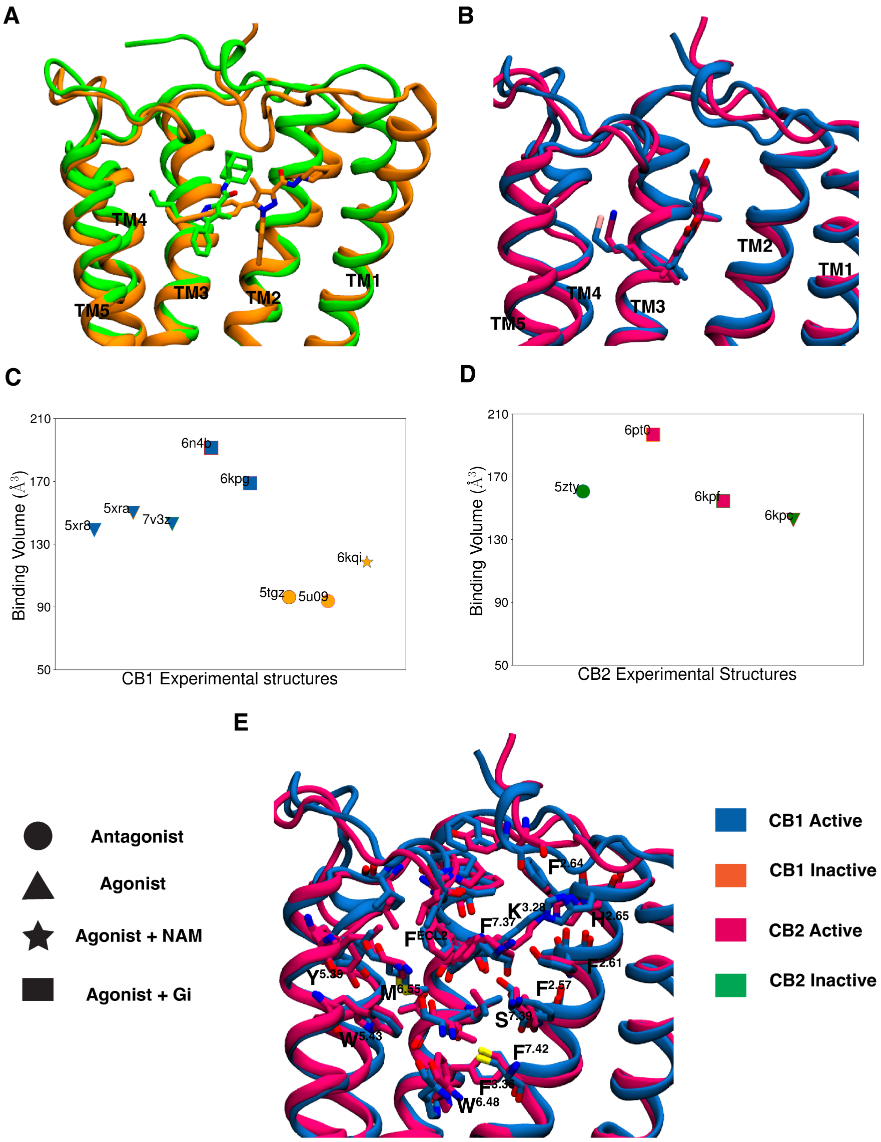


Figure S1: Superposition of inactive (A) and active (B) structures of $CB_1$ (Active PDB ID: 5XRA, color: Orange; Inactive PDB ID: 5TGZ, color: Green) and $CB_2$ (Active PDB ID: 6KPF, color: Purple; Inactive PDB ID: 5ZTY, color: Blue). Proteins are shown as cartoon representation. Agonists ($CB_1$ agonist: AM11542; $CB_2$ agonist: AM841) antagonists ($CB_1$ antagonist: AM6538; $CB_2$ antagonist: AM10257) are shown as sticks. Cartoon representations of TM5 and TM6 are not shown for better visualization of ligand bound positions. Binding pocket volumes for experimentally determined $CB_1$ (C) and $CB_2$ (D) structures are shown as scatter plots. Markers are colored based on the activation state of the protein. The shape of the marker are based on the type of the ligand and downstream signaling partner. Superposition of active $CB_1$ (PDB ID: 5XRA, color: blue) and $CB_2$ (PDB ID: 6KPF, color: purple) structures. Protein structures are shown as cartoon representation. Binding pocket residues are shown as sticks. Cartoon representation of TM6 and TM7 are not shown for better visualization.

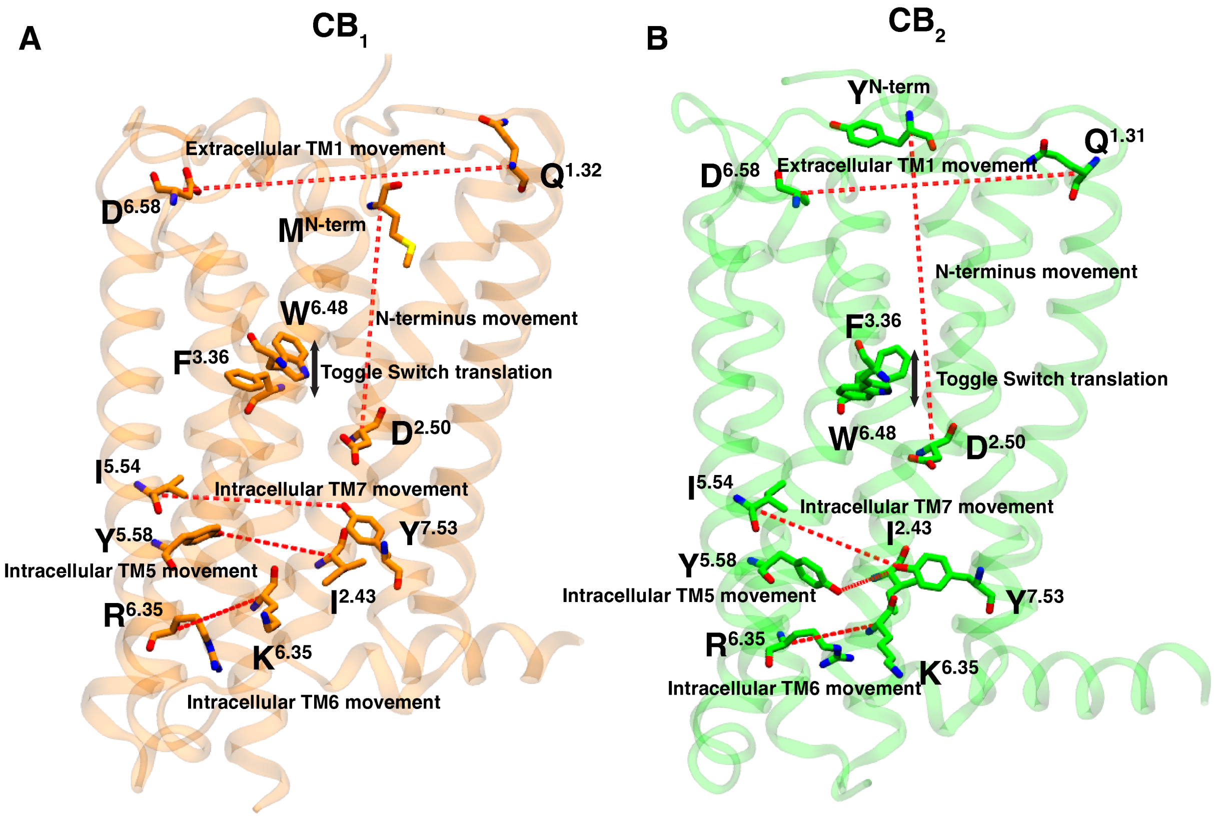


Figure S2: Inactive crystal structure of $CB_1$ (A) (PDB ID: 5TGZ, color: Orange) and $CB_2$ (B) (PDB ID: 5TZY, color: Green) are shown as cartoon representations. Distance metrics used to calculate conserved and non-conserved structural differences between active and inactive states are shown red dotted lines. Toggle switch movements captured by z component difference between $W^{6.48}$ and $F^{3.36}$ are represented as bidirectional arrow.

**Table S1: $CB_1$ and $CB_2$ X-ray crystal and cryo-EM structures bound with different ligands and downstream proteins**

| | **Antagonist** | **Agonist** | **Agonist and NAM** | **Agonist and $G_i$** |
|---|---|---|---|---|
| $CB_1$ | 5TGZ (Inactive)[1]<br>5U09 (Inactive)[5] | 5XRA (Active)[2]<br>5XR8 (Active)[2]<br>7V3Z (Active)[7] | 6KQI (Inactive)[3] | 6N4B (Active)[4]<br>6KPG (Active)[6] |
| $CB_2$ | 5ZTY (Inactive)[8] | 6KPC (Inactive)[6] | | 6KPF (Active)[6]<br>6PT0 (Active)[9] |

**Table S2: Details of apo and holo (agonist and antagonist) unbiased simulation of $CB_1$ and $CB_2$.**

| Protein | Presence of Ligands | Total Simulation time ($\mu$s) |
|---|---|---|
| $CB_1$ | apo (without ligand) | $\sim$ 419 |
| $CB_1$ | holo (with agonist bound) | $\sim$ 24 |
| $CB_1$ | holo (with antagonist bound) | $\sim$ 23 |
| $CB_2$ | apo (without ligand) | $\sim$ 278 |
| $CB_2$ | holo (with agonist bound) | $\sim$ 19 |
| $CB_2$ | holo (with antagonist bound) | $\sim$ 24 |

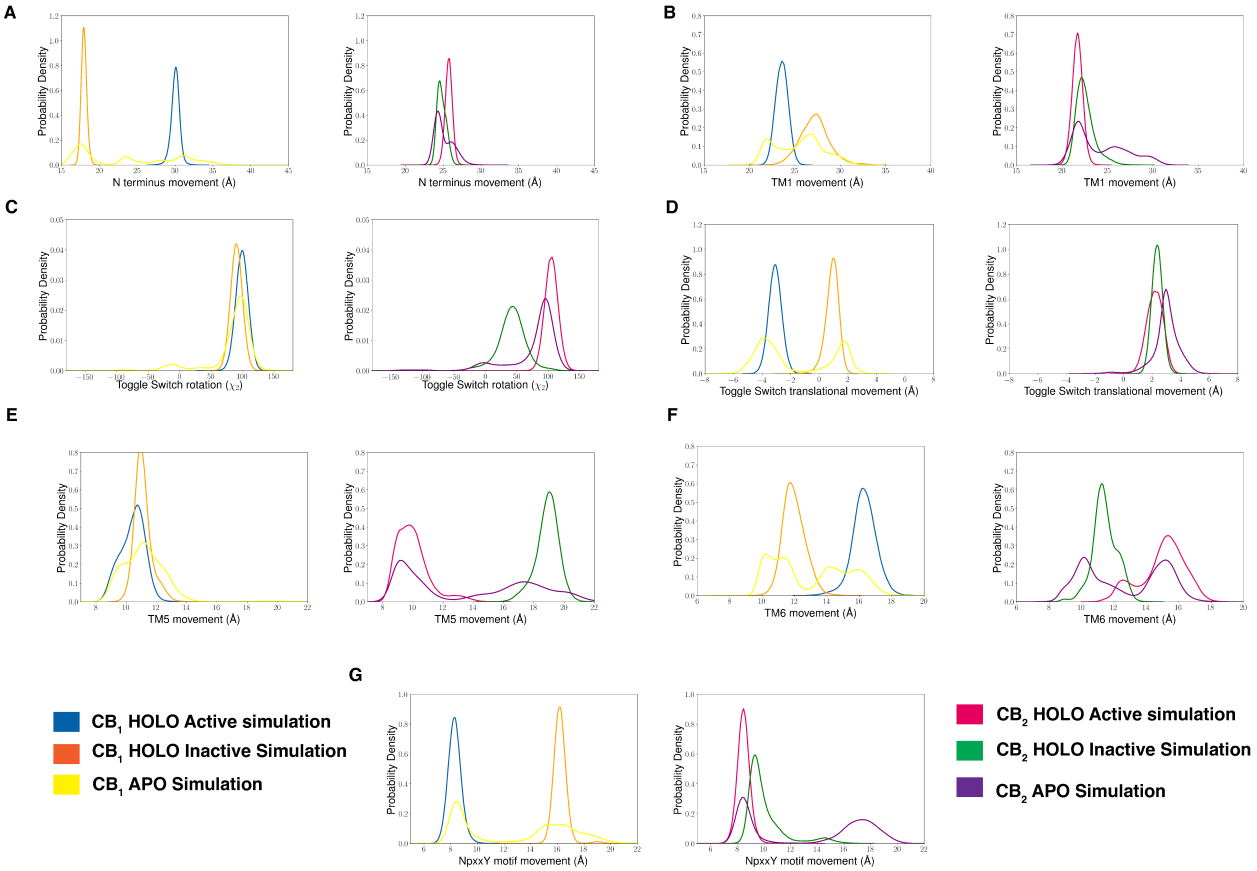


Figure S3: Comparison of distribution of unconserved (A, B, C, D, E) and conserved (F, G) changes of $CB_1$ and $CB_2$ structural features obtained from apo (without ligand) and holo (agonist and antagonist bound) simulation.

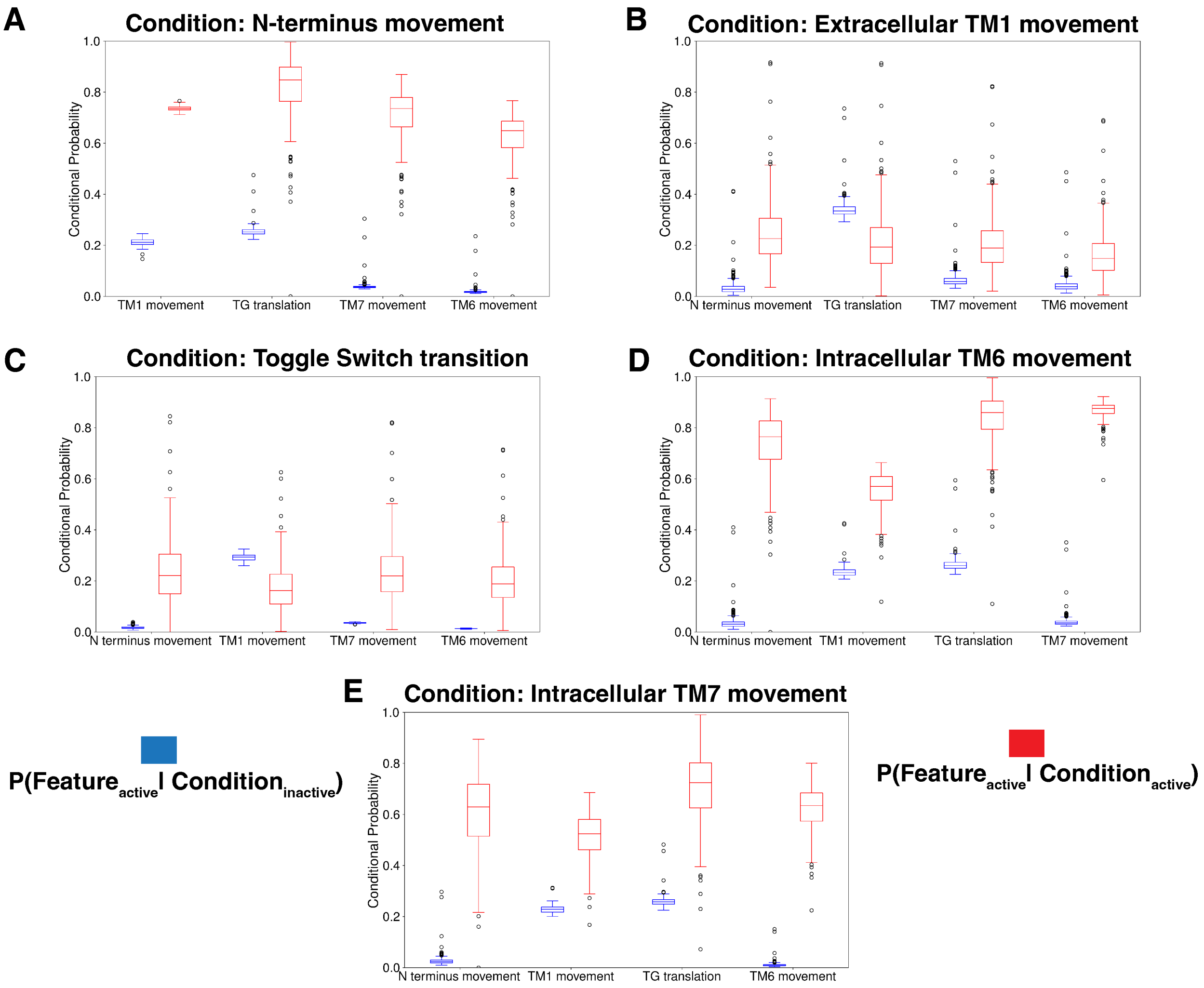


Figure S4: Box plot to show the conditional probabilities for all combinations of structurally important features for $CB_1$. Error calculations were performed based on bootstrapping. 200 bootstrap samples were selected with 80% of the total number of trajectories.

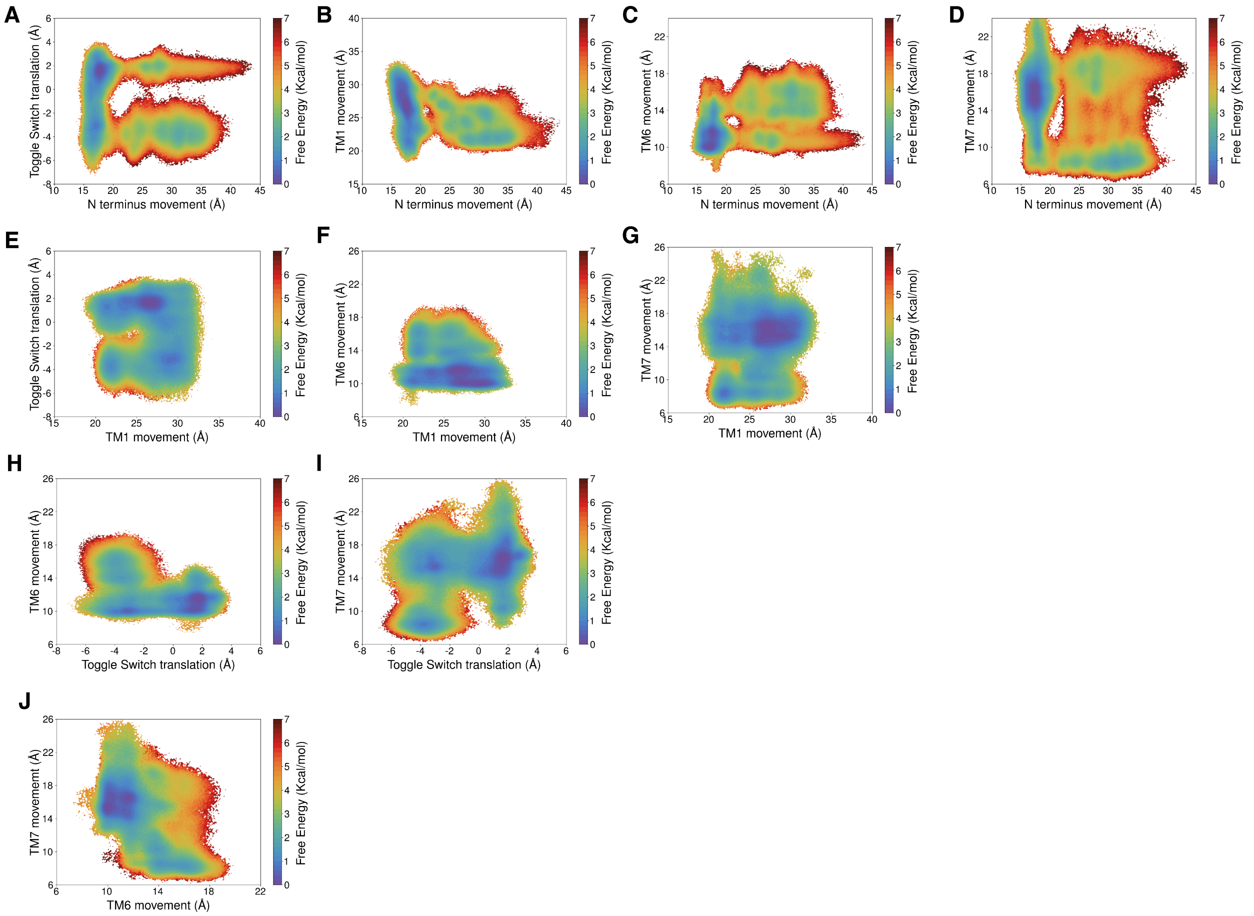


Figure S5: MSM weighted free energy landscapes for all the combinations of dynamically important features for $CB_1$ as discussed in main text.

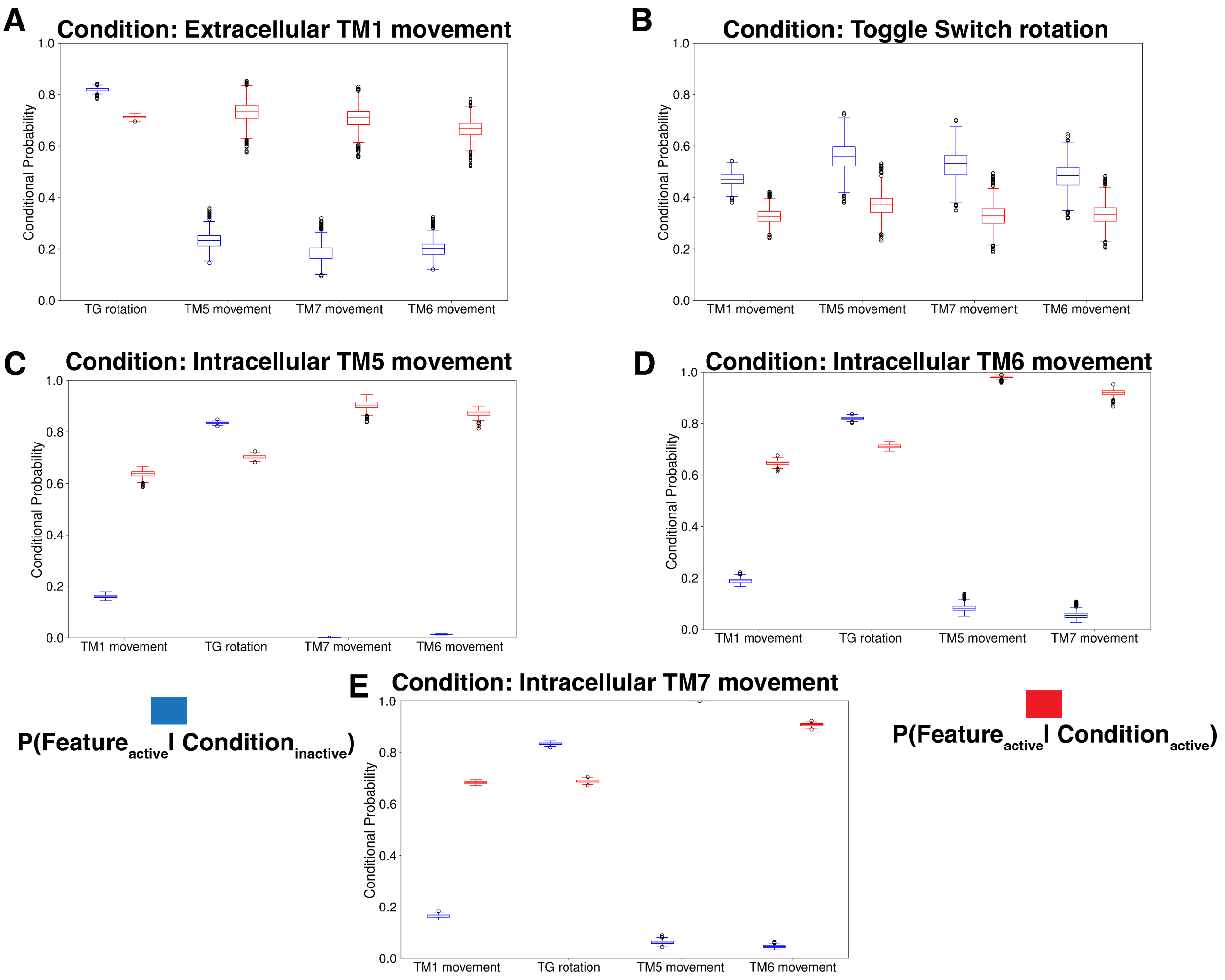


Figure S6: Box plot to show the conditional probabilities for all combinations of structurally important features for $CB_2$. Error calculations were performed based on bootstrapping. 200 bootstrap samples were selected with 80% of the total number of trajectories.

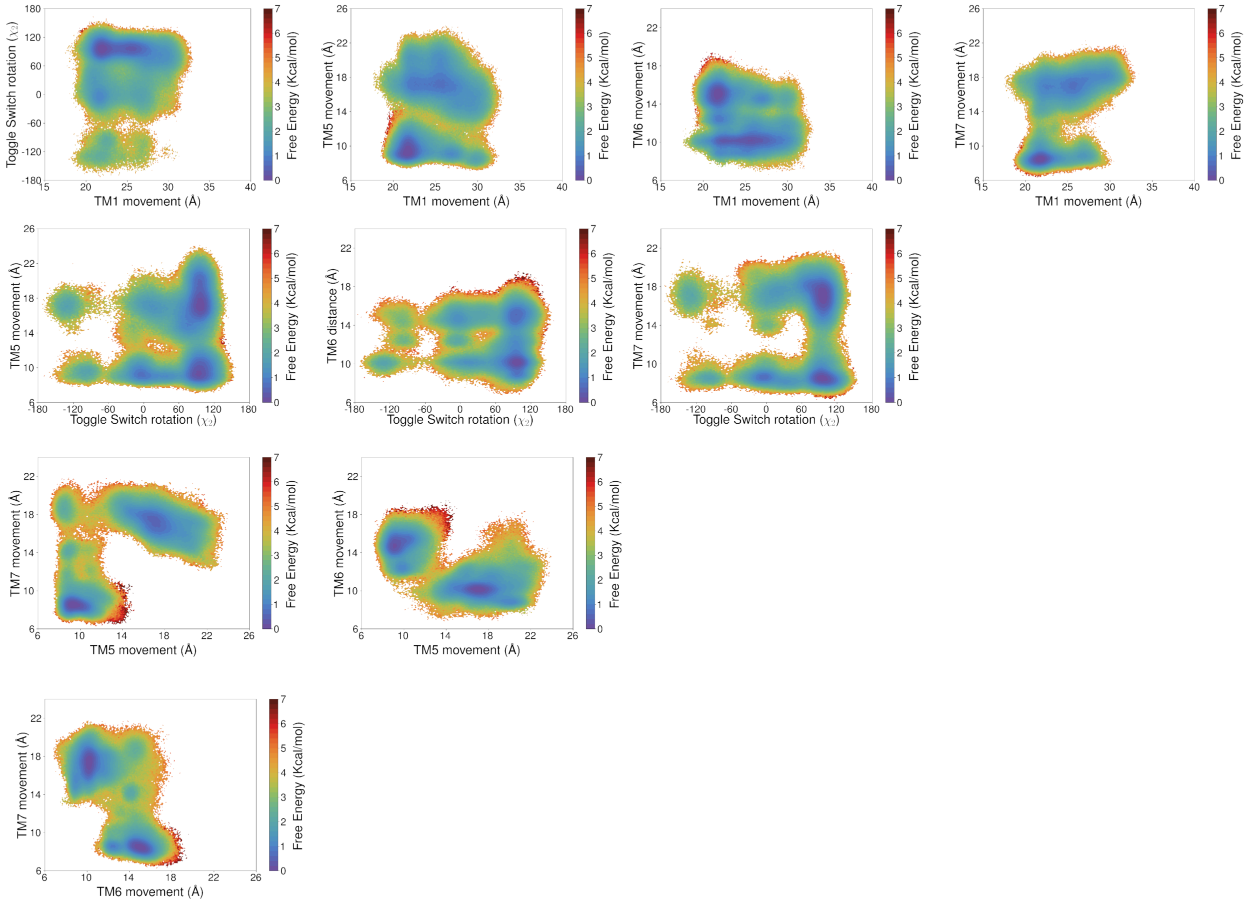


Figure S7: MSM weighted free energy landscapes for all the combinations of dynamically important features as discussed for $CB_2$ in main text.

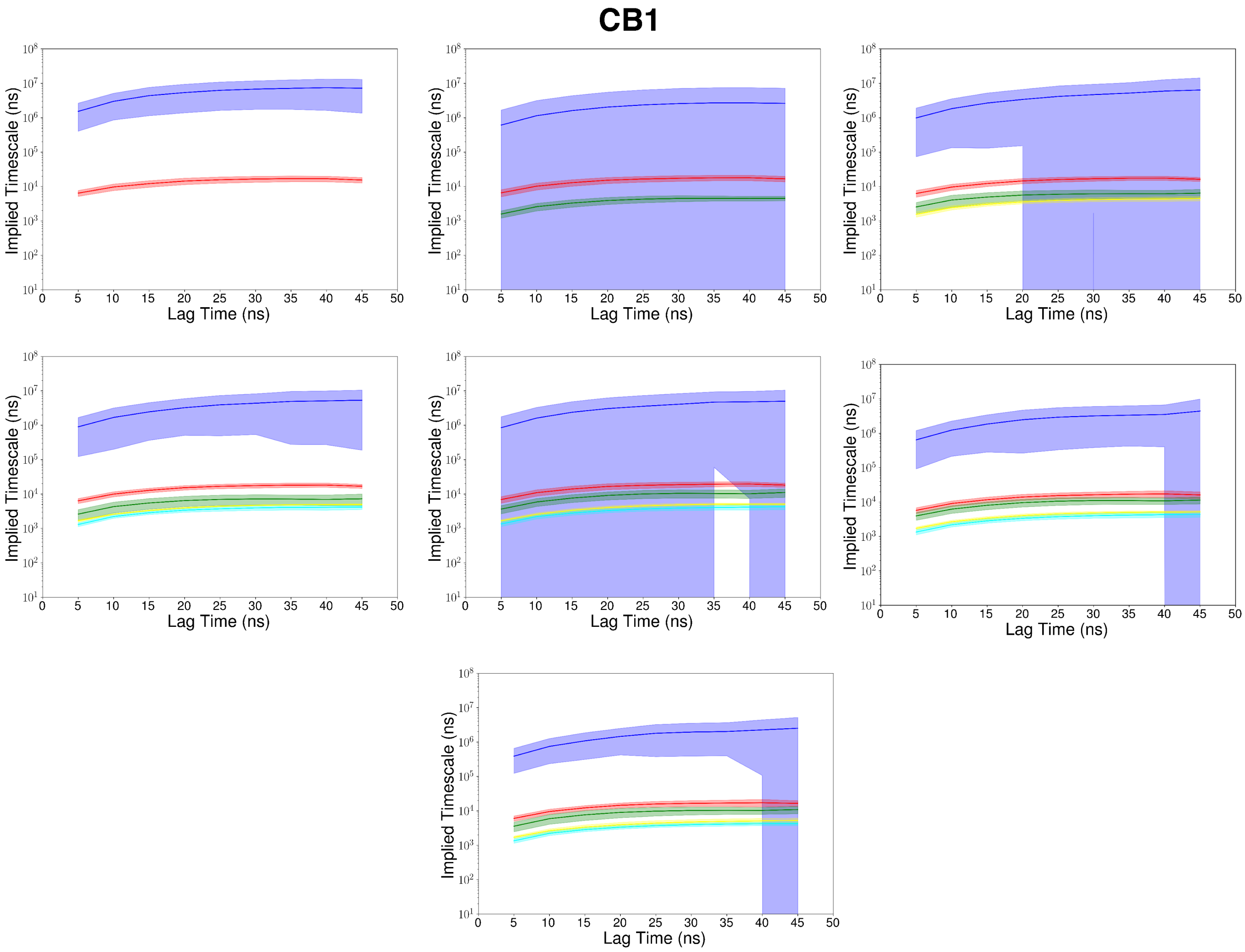


Figure S8: For $CB_1$, implied timecale of VAMPnet models are plotted against the lag time based on the number of the macrostates selected to build the model. Error bars are calculated based on bootstrapping. 20 bootstrap samples were selected with 80% of the total number of trajectories.

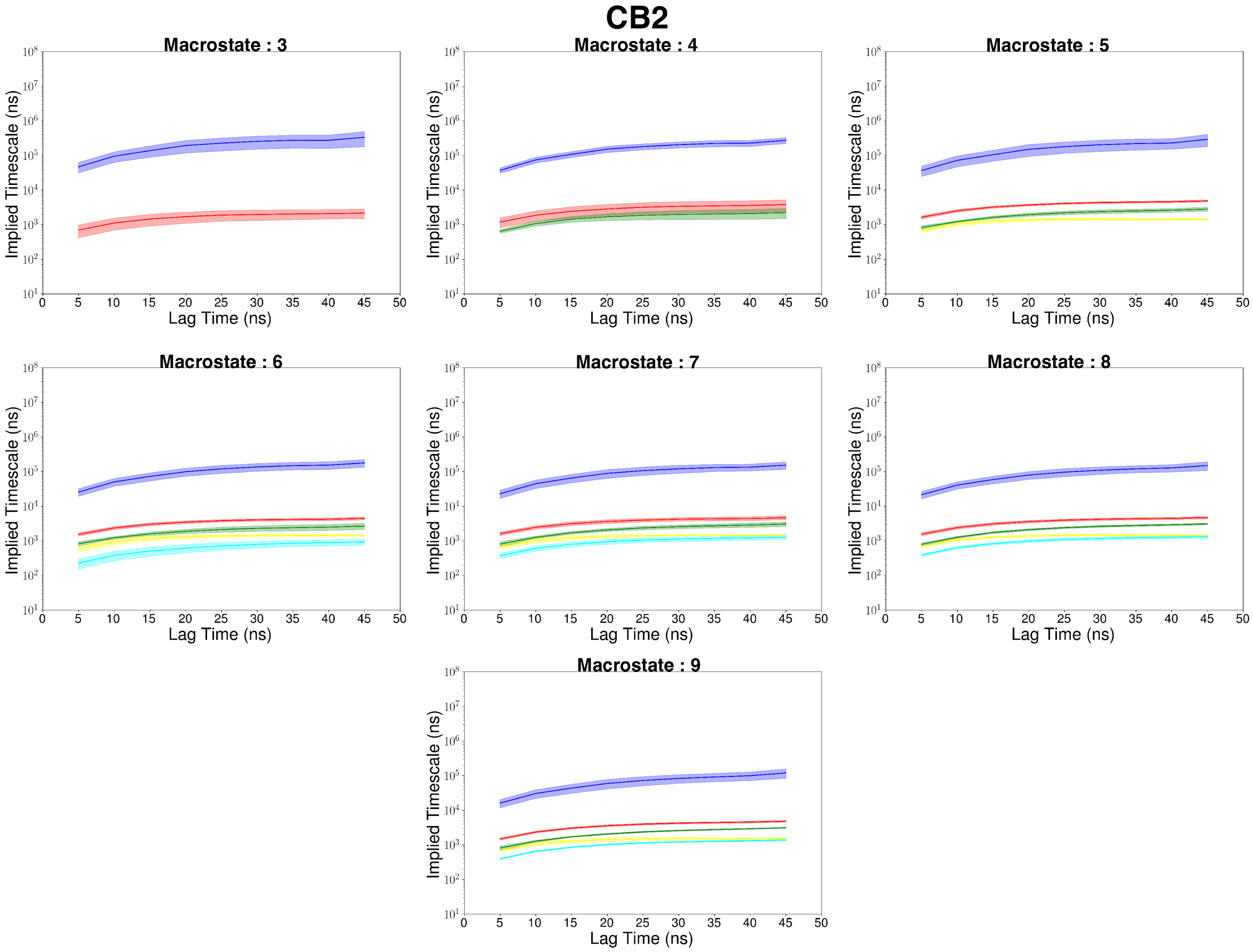


Figure S9: For $CB_2$, implied timecale of VAMPnet models are plotted against the lag time based on the number of the macrostates selected to build the model. Error calculations were performed based on bootstrapping. 20 bootstrap samples were selected with 80% of the total number of trajectories.

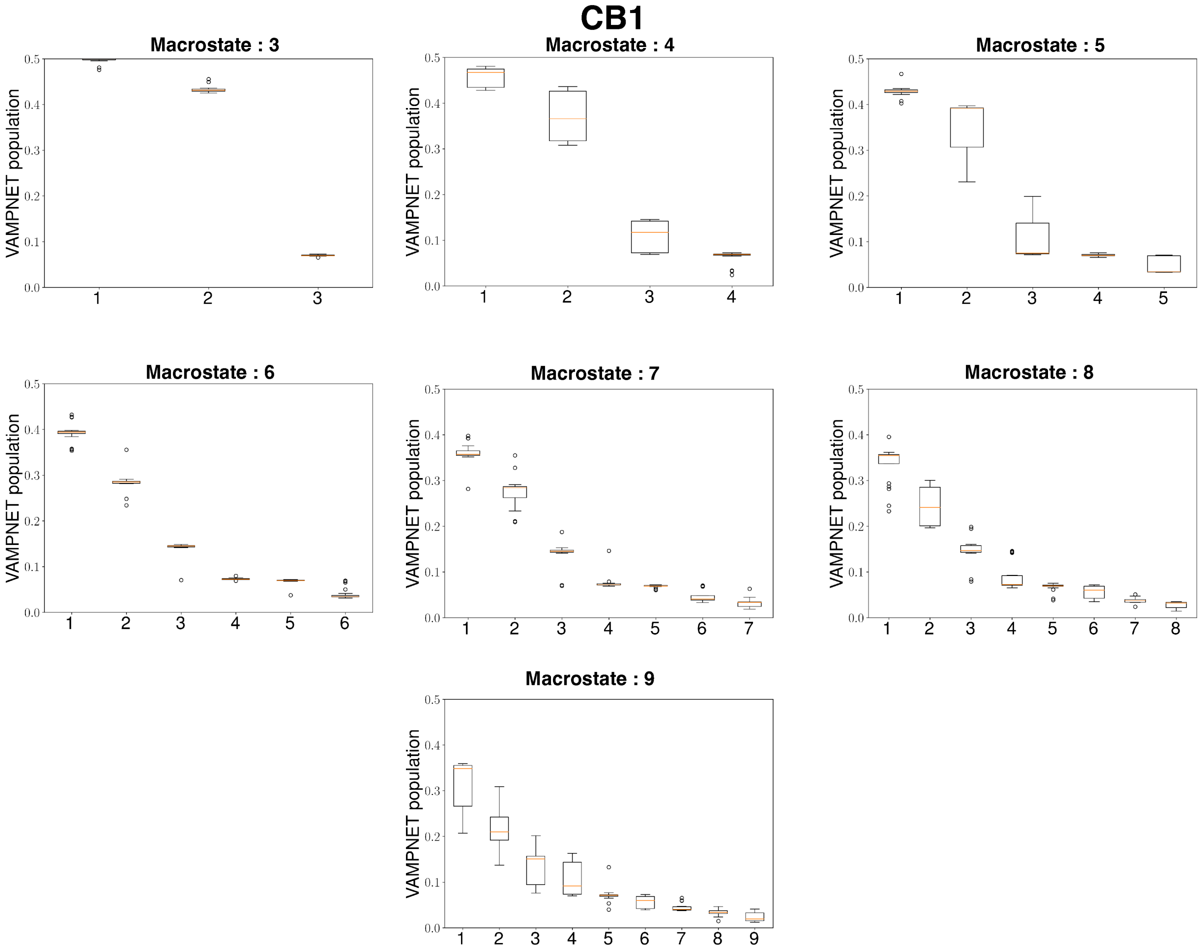


Figure S10: For $CB_1$, state population of VAMPnet models are shown as box plots based on the number of the macrostates selected to build the model. Error calculations were performed based on bootstrapping. 20 bootstrap samples were selected with 80% of the total number of trajectories.

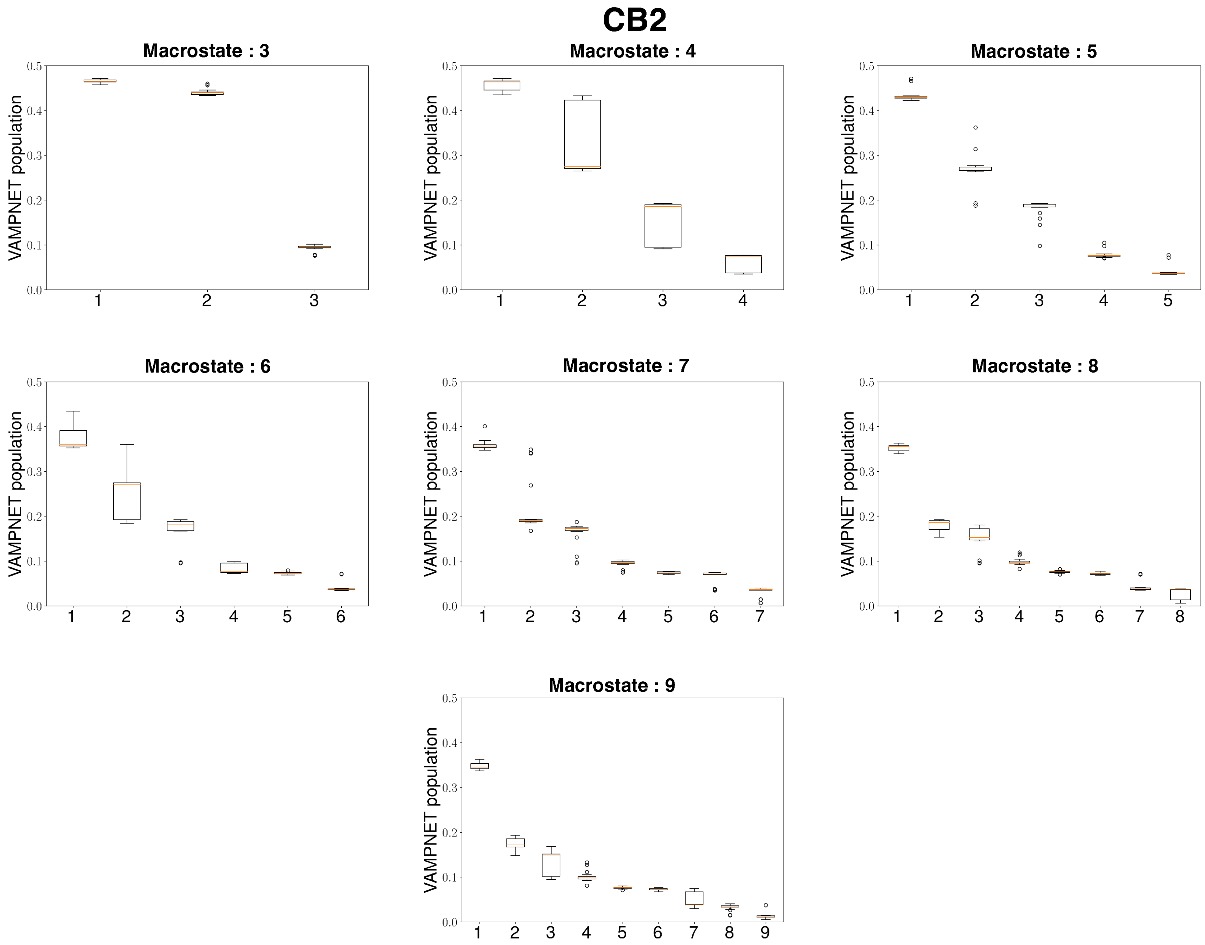


Figure S11: For $CB_2$, state population of VAMPnet models are shown as box plots based on the number of the macrostates selected to build the model. Error bars are calculated based on bootstrapping. 20 bootstrap samples were selected with 80% of the total number of trajectories.

**Table S3: Mean and standard deviation of structurally important features for every metastable state of $CB_1$**

| | Inactive | I1 | I2 | I3 | I4 | Active |
|---|---|---|---|---|---|---|
| N-terminus motion (Å) | $18.35 \pm 0.49$ | $17.12 \pm 0.16$ | $27.91 \pm 1.21$ | $20.12 \pm 0.84$ | $23.43 \pm 0.18$ | $30.72 \pm 0.77$ |
| TM1 movement (Å) | $23.72 \pm 0.54$ | $27.88 \pm 0.37$ | $24.28 \pm 0.22$ | $28.51 \pm 0.25$ | $26.96 \pm 0.1$ | $23.29 \pm 0.39$ |
| TG relative motion (Å) | $1.35 \pm 0.13$ | $-0.09 \pm 0.47$ | $1.9 \pm 0.09$ | $-3.8 \pm 0.13$ | $-4.57 \pm 0.12$ | $-3.66 \pm 0.12$ |
| TM6 movement (Å) | $11.01 \pm 0.21$ | $11.05 \pm 0.2$ | $10.78 \pm 0.09$ | $14.58 \pm 0.16$ | $15.26 \pm 0.22$ | $15.03 \pm 0.21$ |
| TM7 movement (Å) | $16.43 \pm 0.48$ | $15.81 \pm 0.46$ | $17.91 \pm 0.24$ | $9.37 \pm 0.89$ | $8.51 \pm 0.24$ | $9.77 \pm 1.11$ |

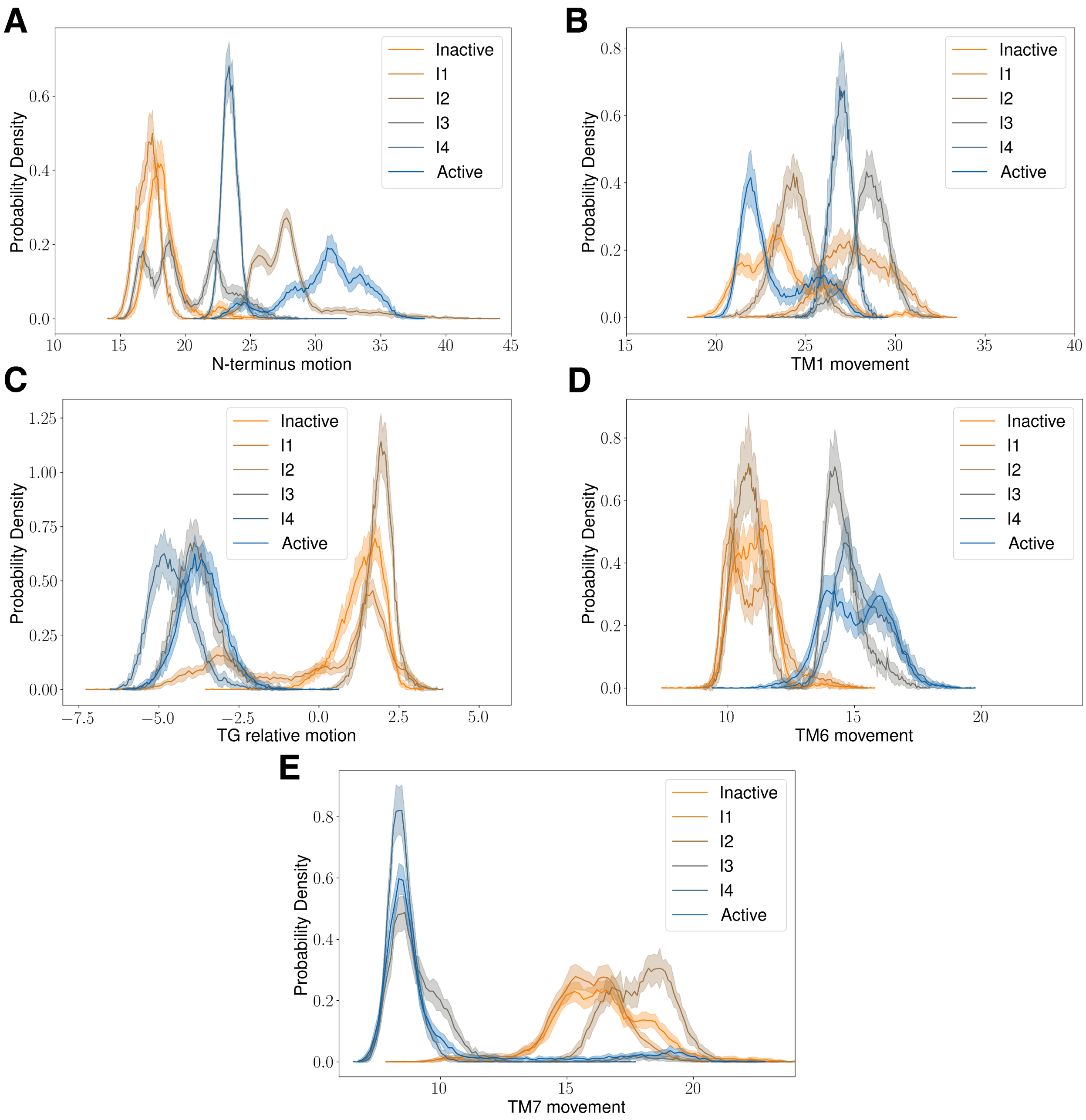


Figure S12: Distributions of structurally important features for each metastable state of $CB_1$.

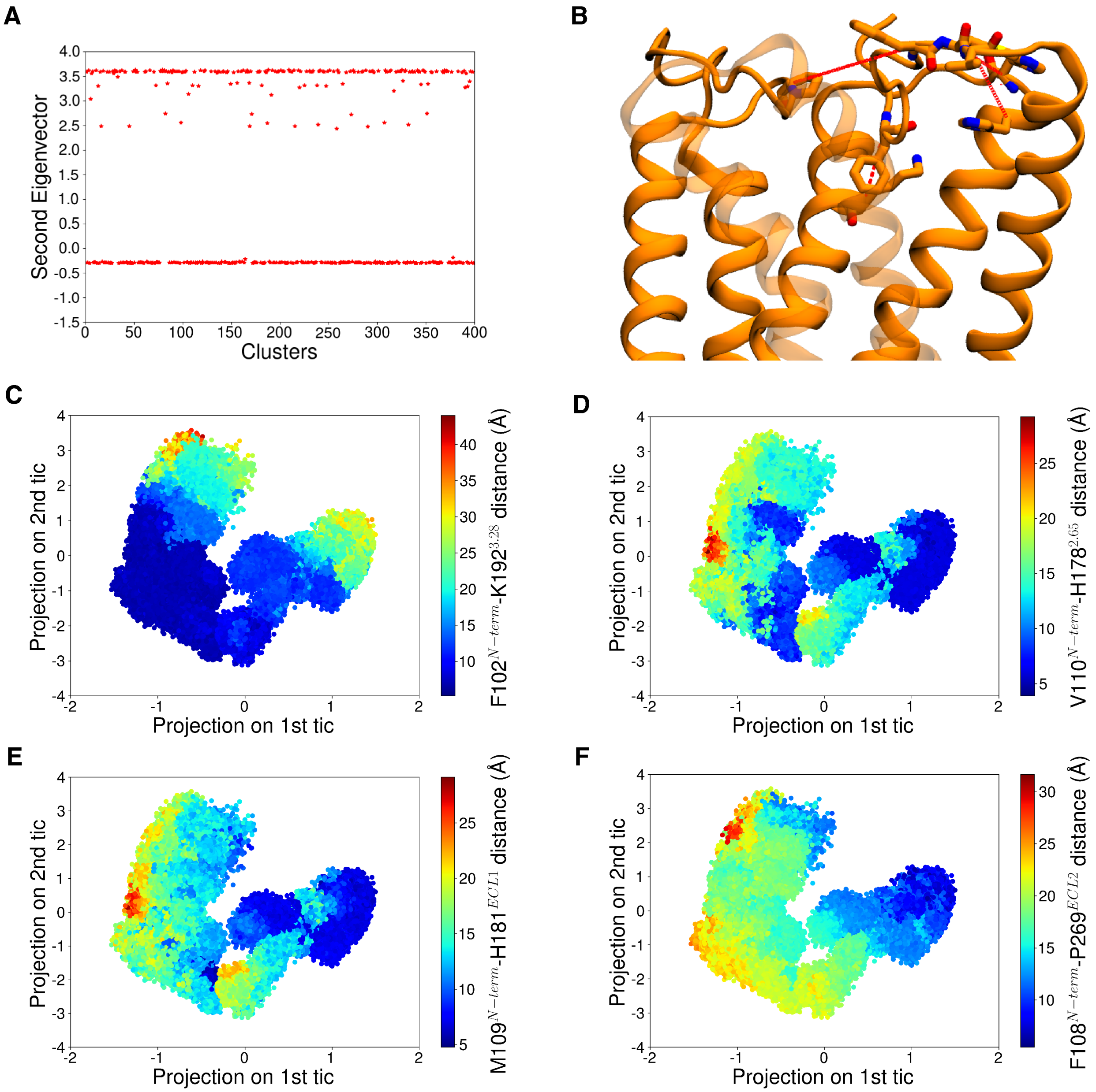


Figure S13: (A) Values of the second eigenvectors of $CB_1$ MSM are plotted as a scatter plot against the cluster numbers. (B) Features that are correlated with the second eigenvectors are calculated are shown as red dotted line. (C, D, E, F) Scatter plots of tic 1 and tic 2 projections of $CB_1$ which are colored based on the values of four features that are highly correlated with second eigenvectors.

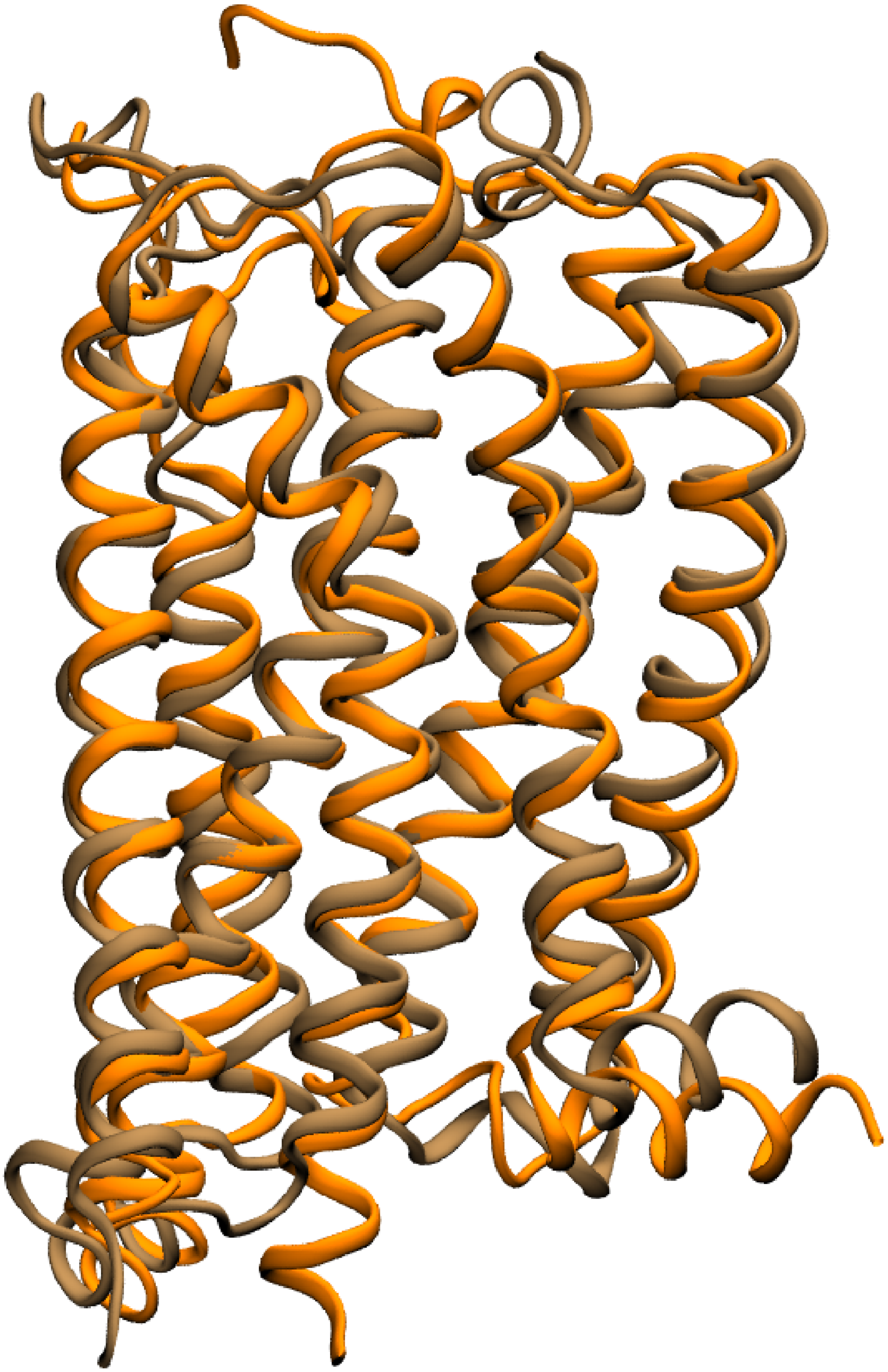

Figure S14: Structural comparison of agonist and NAM bound PDB structure (6KQI, color: Orange) and representative structure from $CB_1$ I2 metastable state.

**Table S4: Mean and standard deviation of structurally important features for every metastable state of $CB_2$**

| | Inactive | I1 | I2 | I3 | I4 | Active |
|---|---|---|---|---|---|---|
| TM1 movement (Å) | $26.05 \pm 0.47$ | $27.86 \pm 0.41$ | $24.06 \pm 0.37$ | $25.0 \pm 0.33$ | $21.85 \pm 0.23$ | $22.41 \pm 0.49$ |
| TG rotation (Å) | $71.97 \pm 17.43$ | $70.3 \pm 9.15$ | $76.04 \pm 24.16$ | $92.52 \pm 9.99$ | $91.79 \pm 12.59$ | $64.97 \pm 11.29$ |
| TM5 movement (Å) | $16.38 \pm 0.79$ | $15.37 \pm 0.28$ | $17.95 \pm 0.29$ | $18.55 \pm 0.41$ | $10.14 \pm 0.36$ | $9.7 \pm 0.21$ |
| TM6 movement (Å) | $10.42 \pm 0.43$ | $9.8 \pm 0.19$ | $9.83 \pm 0.12$ | $10.93 \pm 0.33$ | $15.22 \pm 0.28$ | $14.73 \pm 0.24$ |
| TM7 movement (Å) | $13.73 \pm 0.41$ | $14.85 \pm 0.38$ | $11.73 \pm 0.42$ | $12.92 \pm 0.37$ | $8.57 \pm 0.14$ | $8.82 \pm 0.5$ |

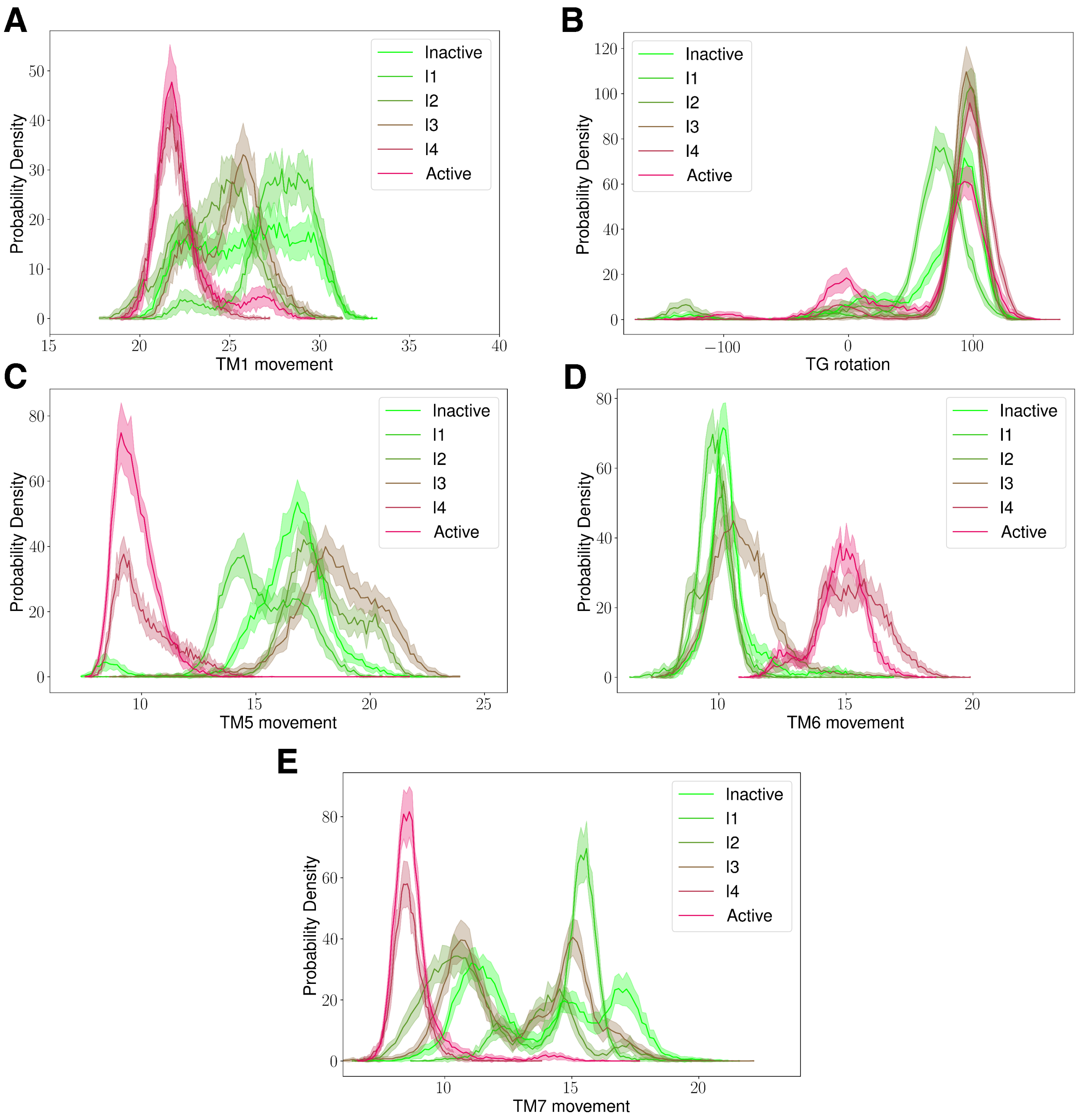


Figure S15: Distributions of structurally important features for each metastable state of $CB_2$.

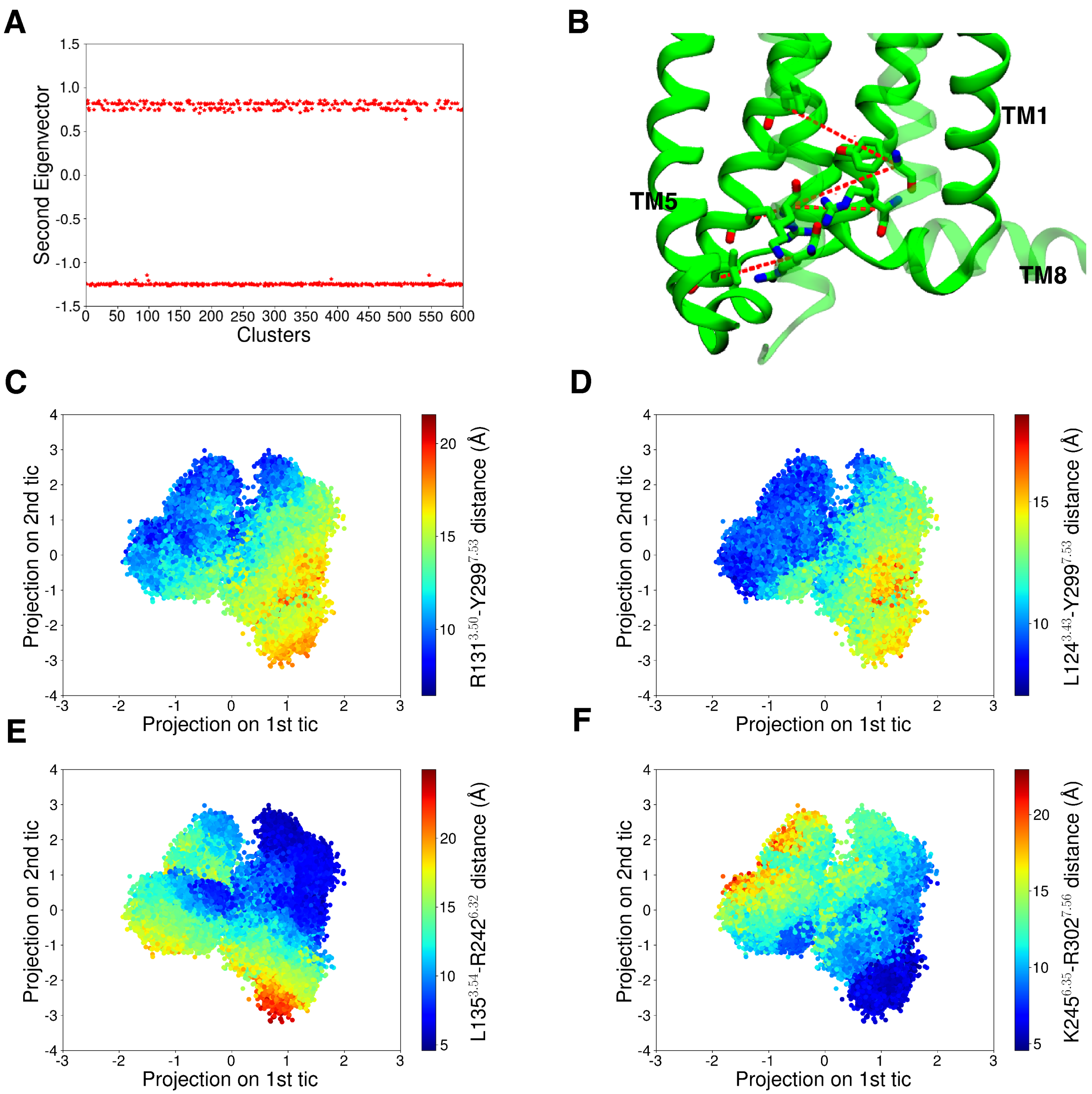


Figure S16: (A) Values of the second eigenvectors of $CB_2$ MSM are plotted as a scatter plot against the cluster numbers. (B) Features that are correlated with the second eigenvectors are calculated are shown as red dotted line. (C, D, E, F) Scatter plots of tic 1 and tic 2 projections of $CB_2$ which are colored based on the values of four features that are highly correlated with second eigenvectors.

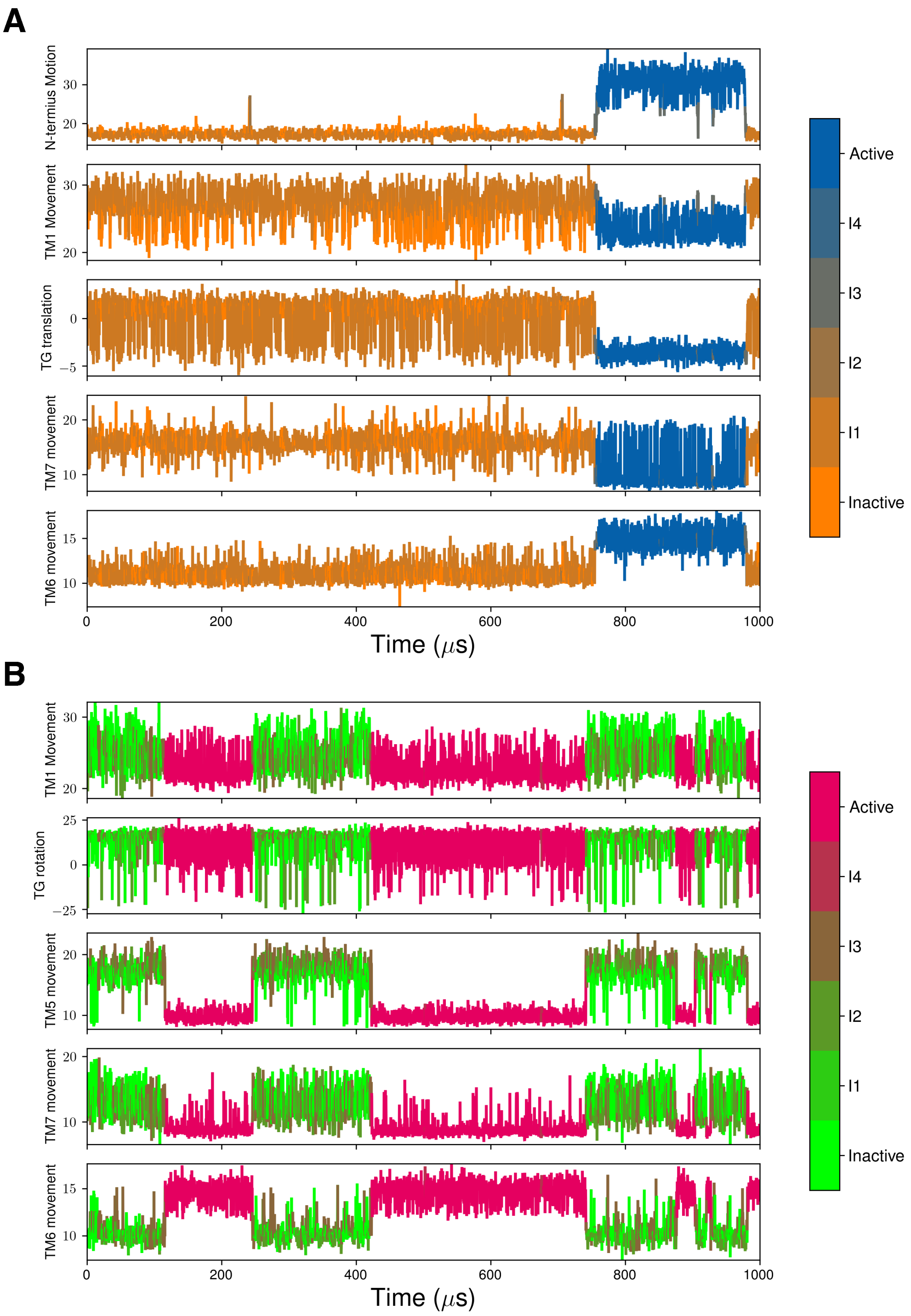


Figure S17: Representative Kinetic monte carlo (kMC) simulation for $CB_1$ and $CB_2$ starting from the inactive state showing the transition of important structural features between different macrostates. Color bar represents different macrostates.

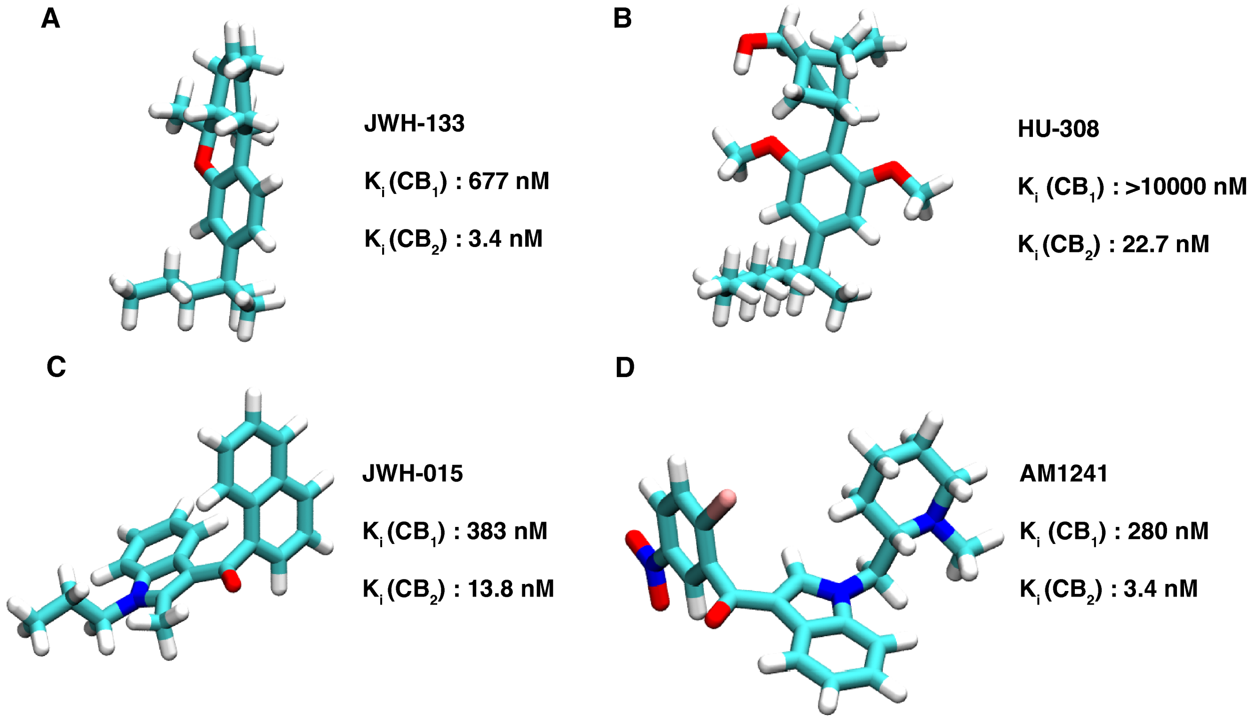


Figure S18: $CB_2$ selective ligands considered for the docking study shown as sticks.

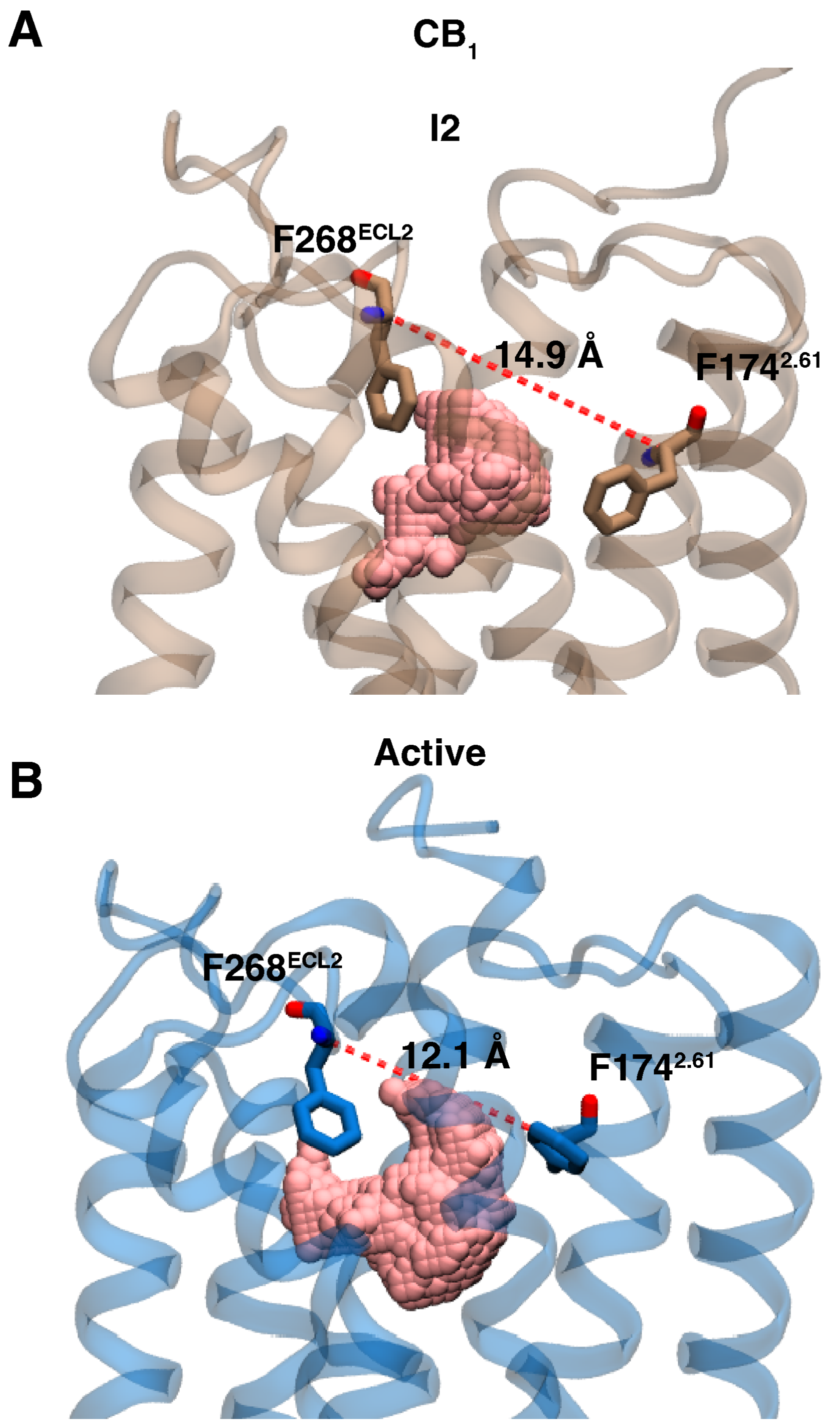


Figure S19: Distance between F268$^{ECL2}$ and F174$^{2.61}$ residues for I2 (A) and active (B) metastable states $CB_1$ are shown as red dotted lines. Proteins are shown as transparent cartoon. Pocket volume calculated using POVME software are shown as beads.

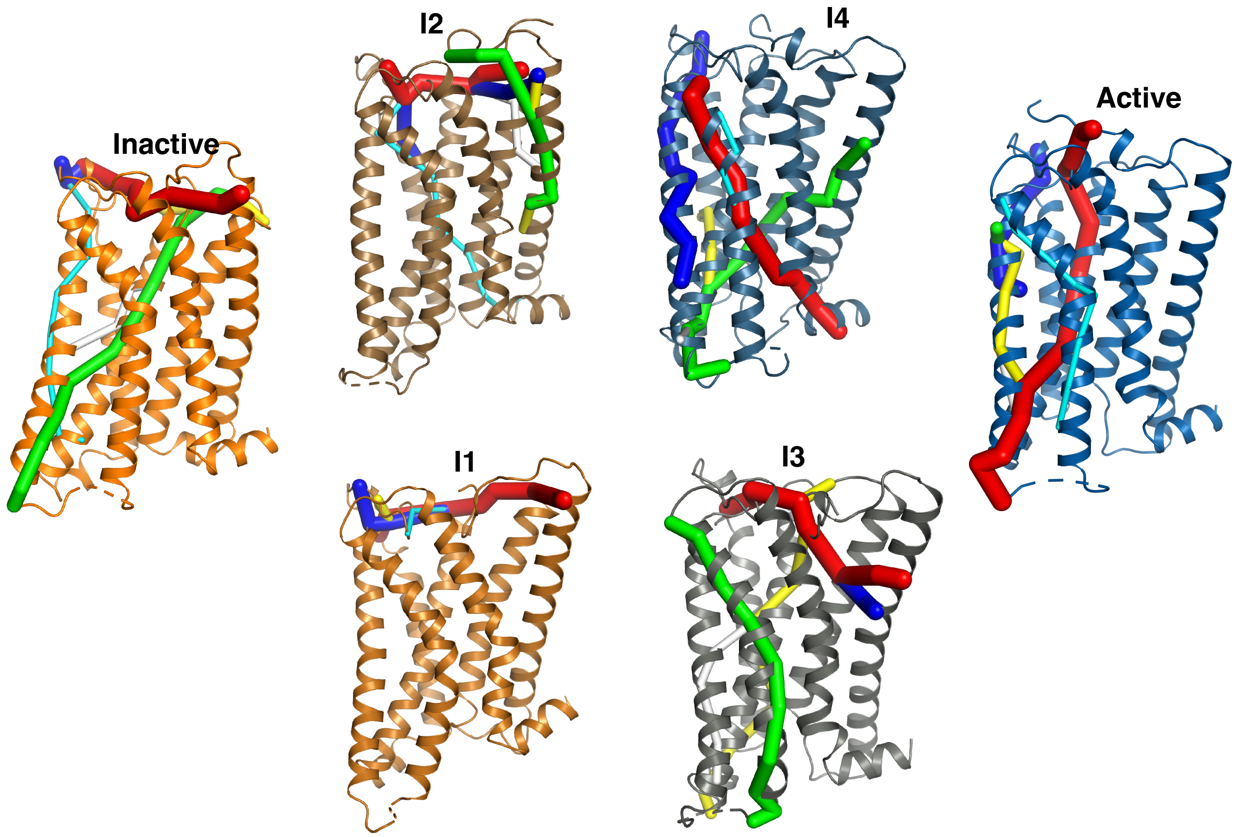


Figure S20: Allosteric communication paths calculated for each metastable state for $CB_1$. Six tunnels with highest numbers of allosteric networks are shown. Tunnel radius are sorted (thicker to thinner) based on the number of allosteric networks in the tunnels.

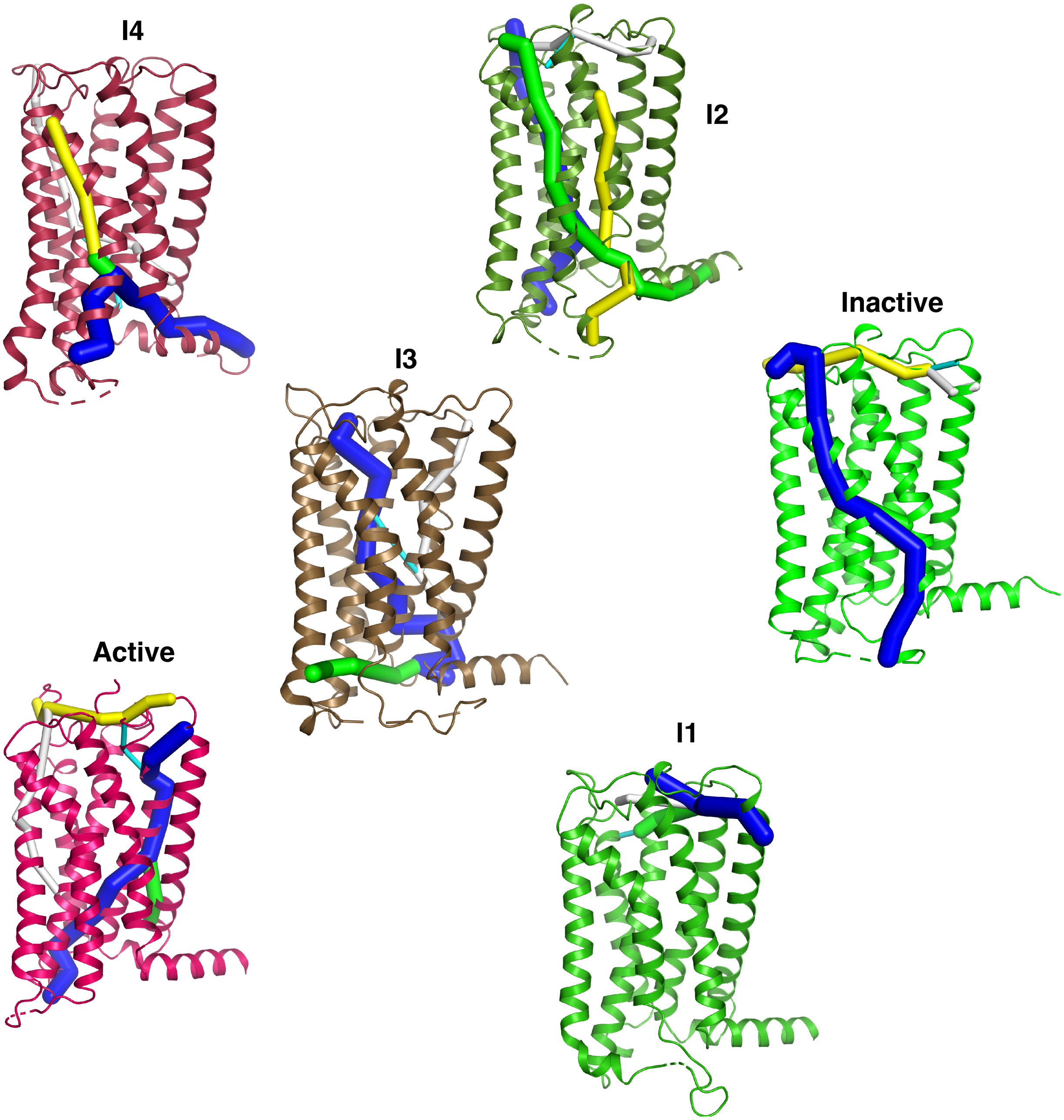


Figure S21: Allosteric communication paths calculated for each metastable state for $CB_2$. Six tunnels with highest numbers of allosteric networks are shown. Tunnel radius are sorted (thicker to thinner) based on the number of allosteric networks in the tunnels.

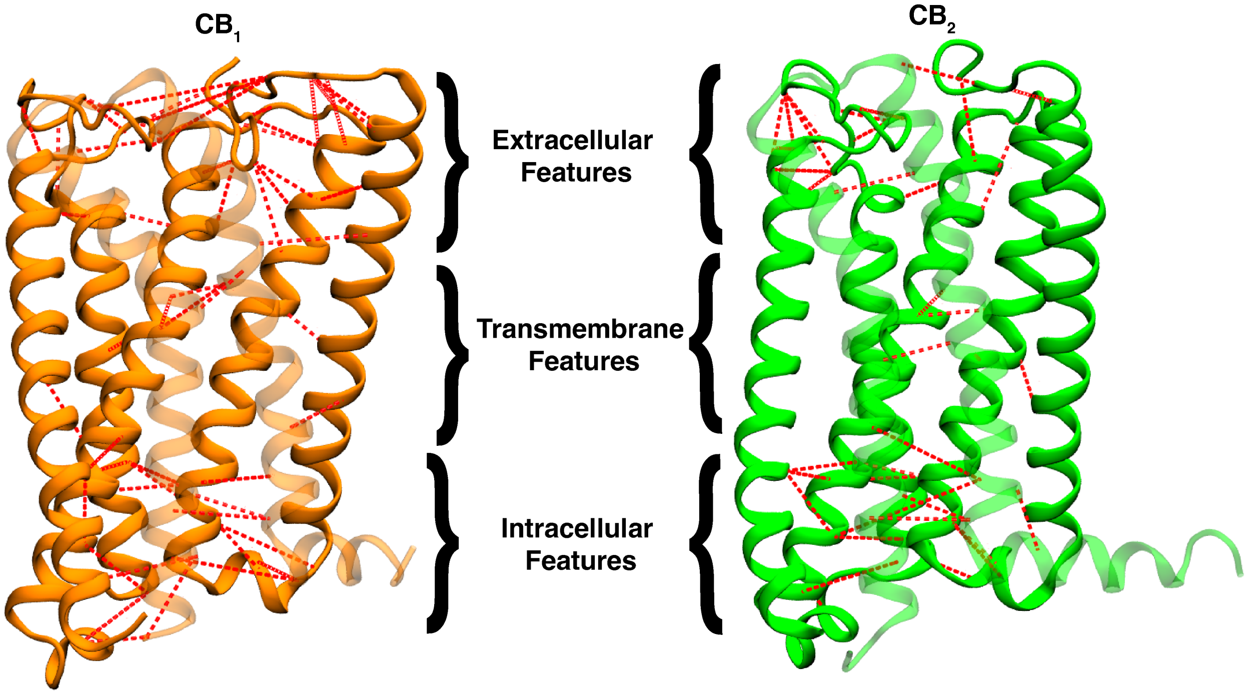


Figure S22: Positions of the features calculated using RRCS analysis are shown as red dotted lines on cartoon representations of CB$_1$ inactive (color: Orange) (A) and CB$_2$ inactive (color: Green) (B) structures.

**Table S5: Distance features used to build Markov state model for $CB_1$ using RRCS.**

| Position | Number | Type | Feature |
|---|---|---|---|
| Extracellular | 1 | $C\alpha$ distance | $E100^{N-term}$-$H270^{ECL2}$ |
| | 2 | | $N101^{N-term}$-$F177^{2.64}$ |
| | 3 | | $N101^{N-term}$-$K183^{ECL1}$ |
| | 4 | | $N101^{N-term}$-$D184^{ECL1}$ |
| | 5 | | $F102^{N-term}$-$F189^{3.25}$ |
| | 6 | | $F102^{N-term}$-$K192^{3.28}$ |
| | 7 | | $M103^{N-term}$-$F170^{2.57}$ |
| | 8 | | $M103^{N-term}$-$S173^{2.60}$ |
| | 9 | | $M103^{N-term}$-$F174^{2.61}$ |
| | 10 | | $D104^{N-term}$-$F177^{2.64}$ |
| | 11 | | $F108^{N-term}$-$D266^{ECL2}$ |
| | 12 | | $F108^{N-term}$-$F268^{ECL2}$ |
| | 13 | | $F108^{N-term}$-$P269^{ECL2}$ |
| | 14 | | $M109^{N-term}$-$F177^{2.64}$ |
| | 15 | | $M109^{N-term}$-$H181^{ECL1}$ |
| | 16 | | $M109^{N-term}$-$Q115^{1.31}$ |
| | 17 | | $M109^{N-term}$-$Q116^{1.32}$ |
| | 18 | | $V110^{N-term}$-$H178^{2.65}$ |
| | 19 | | $A120^{1.36}$-$F174^{2.61}$ |
| | 20 | | $L122^{1.38}$-$M384^{7.40}$ |
| | 21 | | $R186^{3.22}$-$N256^{ECL2}$ |
| | 22 | | $R186^{3.22}$-$E258^{ECL2}$ |
| | 23 | | $F191^{3.27}$-$L252^{4.61}$ |
| | 24 | | $P251^{4.60}$-$Y275^{5.39}$ |

| | | | |
|---|---|---|---|
| | 25 | | $G254^{4.63}$-$K259^{ECL2}$ |
| | 26 | | $E258^{ECL2}$-$H270^{ECL2}$ |
| | 27 | | $S262^{ECL2}$-$E273^{5.37}$ |
| | 28 | | $D266^{ECL2}$-$M371^{ECL3}$ |
| Transmembrane | 29 | | $V131^{1.47}$-$S167^{2.54}$ |
| | 30 | | $E133^{1.49}$-$P394^{7.50}$ |
| | 31 | | $F200^{3.36}$-$W356^{6.48}$ |
| | 32 | | $F200^{3.36}$-$C386^{7.42}$ |
| | 33 | | $T201^{3.37}$-$A244^{4.53}$ |
| | 34 | | $L207^{3.43}$-$Y294^{5.58}$ |
| | 35 | | $F208^{3.44}$-$L286^{5.50}$ |
| | 36 | | $L209^{3.45}$-$F237^{4.46}$ |
| | 37 | | $C355^{6.47}$-$L385^{7.41}$ |
| | 38 | | $W356^{6.48}$-$C386^{7.42}$ |
| Intracellular | 39 | | $S144^{1.60}$-$Y153^{2.40}$ |
| | 40 | | $L147^{ICL1}$-$Y153^{2.40}$ |
| | 41 | | $R148^{ICL1}$-$D403^{8.49}$ |
| | 42 | | $P151^{2.38}$-$D403^{8.49}$ |
| | 43 | | $H154^{2.41}$-$F237^{4.46}$ |
| | 44 | | $F155^{2.42}$-$T210^{3.46}$ |
| | 45 | | $F155^{2.42}$-$Y397^{7.53}$ |
| | 46 | | $F155^{2.42}$-$F237^{4.46}$ |
| | 47 | | $R214^{3.50}$-$Y294^{5.58}$ |
| | 48 | | $R214^{3.50}$-$D338^{6.30}$ |
| | 49 | | $A301^{5.65}$-$R340^{6.33}$ |

| | | | |
|---|---|---|---|
| | 50 | | H304$^{5.68}$-A335$^{6.27}$ |
| | 51 | | H304$^{5.68}$-D338$^{6.30}$ |
| | 52 | | A335$^{6.27}$-R340$^{6.32}$ |
| | 53 | | K343$^{6.35}$-R400$^{7.56}$ |
| | 54 | | T344$^{6.36}$-R400$^{7.56}$ |

**Table S6: Distance features used to build Markov state model for $CB_2$ using RRCS.**

| Position | Number | Type | Feature |
|---|---|---|---|
| Extracellular | 1 | Cα distance | $M22^{N-term}$-$F106^{3.25}$ |
| | 2 | | $I27^{N-term}$-$D275^{7.29}$ |
| | 3 | | $L28^{N-term}$-$H98^{ECL1}$ |
| | 4 | | $K33^{1.32}$-$V96^{2.66}$ |
| | 5 | | $W172^{ECL2}$-$Y190^{5.39}$ |
| | 6 | | $W172^{ECL2}$-$R177^{ECL2}$ |
| | 7 | | $W172^{ECL2}$-$D189^{5.38}$ |
| | 8 | | $T173^{ECL2}$-$L285^{ECL2}$ |
| | 9 | | $C174^{ECL2}$-$L285^{ECL2}$ |
| | 10 | | $R177^{ECL2}$-$P187^{5.36}$ |
| | 11 | | $R177^{ECL2}$-$N188^{5.37}$ |
| | 12 | | $R177^{ECL2}$-$D189^{5.38}$ |
| | 13 | | $E181^{ECL2}$-$K278^{7.32}$ |
| | 14 | | $N188^{5.37}$-$L269^{6.59}$ |
| | 15 | | $D101^{ECL1}$-$K109^{3.28}$ |
| | 16 | | $L107^{3.26}$-$L169^{4.61}$ |
| | 17 | | $L264^{6.54}$-$F281^{7.35}$ |
| Transmembrane | 18 | | $N51^{1.50}$-$F81^{2.51}$ |
| | 19 | | $D80^{2.50}$-$S292^{7.46}$ |
| | 20 | | $A83^{2.53}$-$F117^{3.36}$ |
| | 21 | | $F117^{3.36}$-$C288^{7.42}$ |
| | 22 | | $L124^{3.43}$-$Y299^{7.53}$ |
| | 23 | | $Y209^{5.58}$-$G248^{6.38}$ |
| | 24 | | $L254^{6.44}$-$N291^{7.45}$ |

| | | | |
|---|---|---|---|
| Intracellular | 25 | | L64$^{ICL1}$-Y70$^{2.40}$ |
| | 26 | | R66$^{ICL1}$-R147$^{4.39}$ |
| | 27 | | Y70$^{2.40}$-E305$^{8.49}$ |
| | 28 | | F72$^{2.42}$-L126$^{3.45}$ |
| | 29 | | F72$^{2.42}$-T127$^{3.46}$ |
| | 30 | | A128$^{3.47}$-Y209$^{5.58}$ |
| | 31 | | R131$^{3.50}$-Y209$^{5.58}$ |
| | 32 | | R131$^{3.50}$-Y299$^{7.53}$ |
| | 33 | | R131$^{3.50}$-L243$^{6.33}$ |
| | 34 | | L133$^{3.52}$-Y141$^{ICL2}$ |
| | 35 | | L135$^{3.54}$-R242$^{6.32}$ |
| | 36 | | K245$^{6.35}$-R302$^{7.56}$ |
| | 37 | | T246$^{6.36}$-R302$^{7.56}$ |
| | 38 | | A300$^{7.54}$-R307$^{8.51}$ |

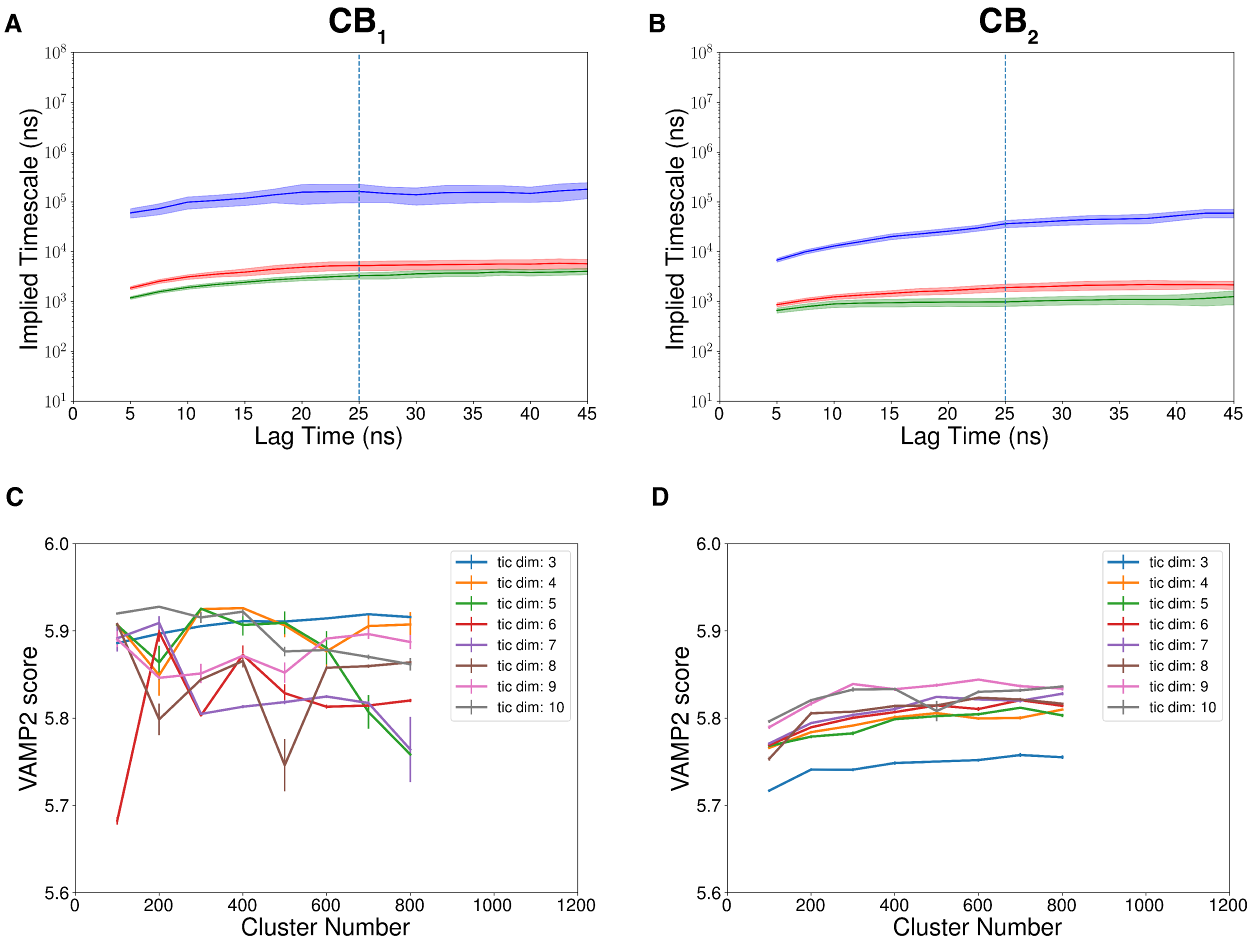


Figure S23: First three implied timescale obtained from MSM is plotted against lag time for $CB_1$ (A) and $CB_2$ (B). Error bars are calculated using bootstrapping method by building 10 different samples with 80% of the total number of trajectories. 25 ns was selected as lagtime to build final MSM. VAMP-2 score was plotted against cluster numbers for $CB_1$ (C) and $CB_2$ (D). These calculations were performed for different tic-dimensions. Error bars were calculated by tenfold cross validation.

**Table S7: Distance and angle features used for adaptive sampling of $CB_1$.**

| Position | Number | Type | Feature |
|---|---|---|---|
| Extracellular helical movement | 1 | distance | $Q^{1.32}(C\alpha)$-$F^{3.25}(C\alpha)$ |
| | 2 | distance | $Q^{1.32}(C\alpha)$-$P^{4.60}(C\alpha)$ |
| | 3 | distance | $Q^{1.32}(C\alpha)$-$L^{5.40}(C\alpha)$ |
| | 4 | distance | $Q^{1.32}(C\alpha)$-$D^{6.58}(C\alpha)$ |
| | 5 | distance | $Q^{1.32}(C\alpha)$-$T^{7.33}(C\alpha)$ |
| | 6 | distance | $D^{2.63}(C\alpha)$-$F^{3.25}(C\alpha)$ |
| | 7 | distance | $D^{2.63}(C\alpha)$-$P^{4.60}(C\alpha)$ |
| | 8 | distance | $D^{2.63}(C\alpha)$-$L^{5.40}(C\alpha)$ |
| | 9 | distance | $D^{2.63}(C\alpha)$-$D^{6.58}(C\alpha)$ |
| | 10 | distance | $D^{2.63}(C\alpha)$-$T^{7.33}(C\alpha)$ |
| N-loop movement | 11 | distance | $M103^{N-term}(C\alpha)$-$D^{2.50}(C\alpha)$ |
| | 12 | distance | $M103^{N-term}(C\alpha)$-$F268^{ECL2}(C\alpha)$ |
| Toggle switch movement | 13 | Dihedral Angle ($\chi_2$) | $F^{3.36}$ |
| | 14 | Dihedral Angle ($\chi_2$) | $W^{6.48}$ |
| | 15 | distance | $W^{6.48}(N\epsilon)$-$D^{2.50}(C\alpha)$ |
| | 16 | distance | $W^{6.48}(N\epsilon)$-$L^{5.50}(C\alpha)$ |
| | 17 | distance | $F^{3.36}(C\gamma)$-$D^{2.50}(C\alpha)$ |
| | 18 | distance | $F^{3.36}(C\gamma)$-$L^{5.50}(C\alpha)$ |
| Intracellular helical movement | 19 | distance | $R^{3.50}(C\alpha)$-$K^{6.35}$ $(C\alpha)$ |
| | 20 | distance | $Y^{2.40}(OH)$-$Y^{7.53}(OH)$ |
| | 21 | distance | $I^{5.54}(C\alpha)$-$Y^{7.53}(OH)$ |
| | 22 | distance | $A^{4.45}(C\alpha)$-$F^{2.42}(C\gamma)$ |
| | 23 | Dihedral Angle ($\chi_2$) | $F^{2.42}$ |
| | 24 | Dihedral Angle ($\chi_2$) | $F^{4.46}$ |

**Table S8: Distance and angle features used for adaptive sampling of $CB_2$.**

| Position | Number | Type | Feature |
|---|---|---|---|
| Extracellular helical movement | 1 | distance | $Q^{1.31}(C\alpha)$-$F^{3.25}(C\alpha)$ |
| | 2 | distance | $Q^{1.31}(C\alpha)$-$P^{4.60}(C\alpha)$ |
| | 3 | distance | $Q^{1.31}(C\alpha)$-$L^{5.40}(C\alpha)$ |
| | 4 | distance | $Q^{1.31}(C\alpha)$-$S^{6.58}(C\alpha)$ |
| | 5 | distance | $Q^{1.31}(C\alpha)$-$K^{7.33}(C\alpha)$ |
| | 6 | distance | $N^{2.63}(C\alpha)$-$F^{3.25}(C\alpha)$ |
| | 7 | distance | $N^{2.63}(C\alpha)$-$P^{4.60}(C\alpha)$ |
| | 8 | distance | $N^{2.63}(C\alpha)$-$L^{5.40}(C\alpha)$ |
| | 9 | distance | $N^{2.63}(C\alpha)$-$D^{6.58}(C\alpha)$ |
| | 10 | distance | $N^{2.63}(C\alpha)$-$K^{7.33}(C\alpha)$ |
| N-loop movement | 11 | distance | $M26^{N-term}(C\alpha)$-$D^{2.50}(C\alpha)$ |
| | 12 | distance | $M26^{N-term}(C\alpha)$-$F183^{ECL2}(C\alpha)$ |
| Toggle switch movement | 13 | Dihedral Angle ($\chi_2$) | $W^{6.48}$ |
| | 14 | distance | $W^{6.48}(N\epsilon)$-$D^{2.50}(C\alpha)$ |
| | 15 | distance | $W^{6.48}(N\epsilon)$-$L^{5.50}(C\alpha)$ |
| | 16 | distance | $F^{3.36}(C\gamma)$-$D^{2.50}(C\alpha)$ |
| | 17 | distance | $F^{3.36}(C\gamma)$-$L^{5.50}(C\alpha)$ |
| Intracellular helical movement | 18 | distance | $R^{3.50}(C\alpha)$-$K^{6.35}$ $(C\alpha)$ |
| | 19 | distance | $Y^{2.40}(OH)$-$Y^{7.53}(OH)$ |
| | 20 | distance | $I^{5.54}(C\alpha)$-$Y^{7.53}(OH)$ |

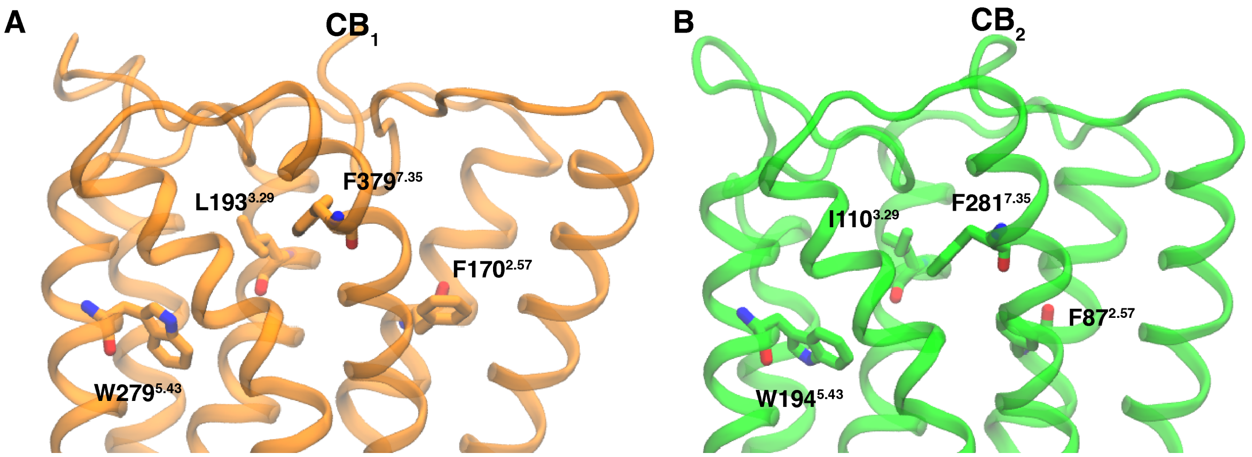


Figure S24: Cartoon representation of $CB_1$ (A) and $CB_2$ (B) showing the binding pocket residues as sticks. Center of mass of the $\alpha$-carbon of these residues were considered as docking center for docking calculation.
